# Failure of Bacillus Calmette-Guérin therapy in patients with bladder cancer is characterized by immune dysfunction associated with Activator Protein 1

**DOI:** 10.64898/2026.05.06.723215

**Authors:** Andrew Garven, Jean-François Paré, Alexandra Robins, Ana Vera-Rodríguez, Rohan Sampy, Arianna Malan-Bennett, Richard W. Nauman, Andrew W. Craig, Peter A. Greer, Madhuri Koti, Tiziana Cotechini, David M. Berman, Amber Simpson, Lynne-Marie Postovit, D. Robert Siemens, Charles H. Graham

**Affiliations:** Department of Biomedical and Molecular Sciences; Department of Public Health Sciences; Department of Pathology and Molecular Medicine; Sinclair Cancer Research Institute and; School of Computing, Queen’s University, Kingston, Ontario, Canada; Department of Radiology & Diagnostic Imaging, University of Alberta, Edmonton, Alberta, Canada; Department of Urology, Queen’s University, Kingston, Ontario, Canada

## Abstract

The standard-of-care for patients with higher-risk non-muscle invasive bladder cancer (NMIBC) after tumour resection is intravesical administration of Bacillus Calmette-Guérin (BCG). While this form of adjuvant immunotherapy has improved recurrence-free and progression-free survival, a large proportion of patients experience recurrences within a year of diagnosis. The reasons for this high rate of early recurrence following BCG therapy remain unclear; however, inadequate activation of systemic immunity may be a contributing factor. To address this, we analysed the transcriptomic and chromatin accessibility profiles of peripheral blood mononuclear cells obtained from patients with NMIBC at single-cell resolution before BCG immunotherapy and after five induction doses of BCG. Monocytes from patients who experienced disease recurrence within a year of initiation of BCG therapy (BCG non-responders) exhibited a pro-inflammatory phenotype consistent with age-related immunosenescence prior to BCG immunotherapy. Moreover, inflammation-associated pathways that were active before initiation of BCG therapy in the BCG non-responders were down-regulated after five instillations of BCG. In contrast, these pathways were quiescent before BCG therapy in patients who remained disease-free for at least a year but were markedly up-regulated after five doses of BCG. Genomic regions with accessible chromatin were enriched in activator protein 1 (AP-1) binding sequences in monocytes from BCG-non-responders prior to BCG therapy. AP-1 is a central regulator of the inflammatory phenotype associated with immunosenescence. Our findings indicate that a pre-existing state of innate immunosenescence underlies early disease recurrence following BCG. Patients unlikely to benefit from BCG may be offered alternative therapies early in their disease journey.

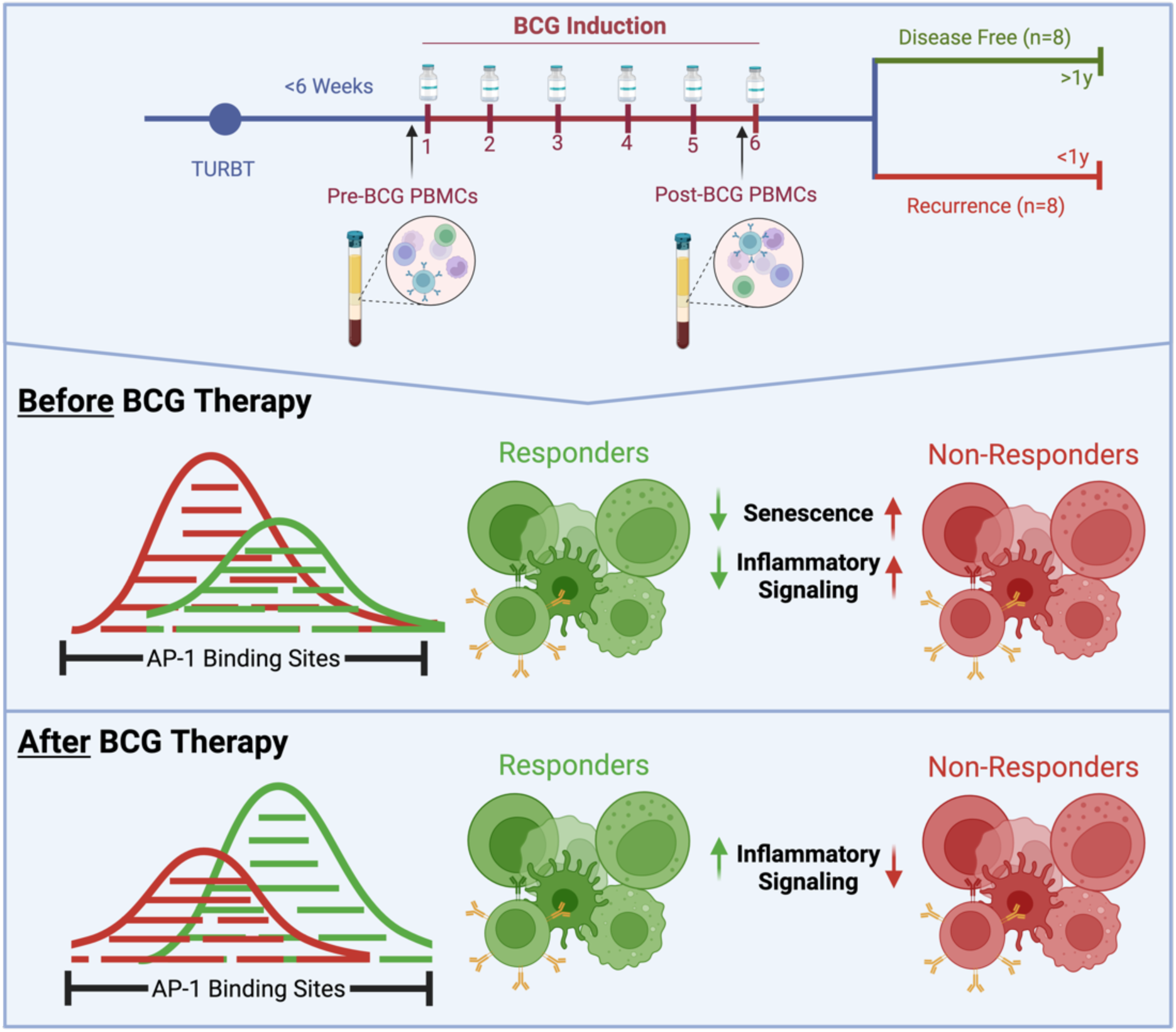

## Introduction

To reduce risk of cancer recurrence and progression, patients with higher risk non-muscle invasive bladder cancer (NMIBC) undergo surgical resection of the tumour followed by intravesical administration of Bacillus Calmette-Guérin (BCG), a live, attenuated form of *Mycobacterium bovis* first used as a vaccine against tuberculosis^1^. Initial BCG treatment involves a six-week induction phase, during which patients receive one dose per week; however, maximal therapeutic efficacy requires one year of maintenance dosing for patients with intermediate-risk and up to three years of maintenance dosing for high-risk patients^2,3^. It is well established that, compared with surgical resection alone, surgical resection followed by BCG immunotherapy increases recurrence-free survival^1,4^.

Despite the proven clinical utility of BCG, up to 61% of patients suffer local disease recurrence within one year of therapy initiation^5,6^, with some (up to 17%) progressing to muscle-invasive cancer^5,7–9^. The reasons for this high incidence of disease recurrence remain unclear; however, one possibility is that patients who experience early disease recurrence following BCG therapy fail to mount effective systemic anti-cancer immune responses. For instance, the inability of peripheral blood monocytes to acquire trained immunity following BCG therapy was associated with an increased incidence of disease recurrence and shorter recurrence-free survival in patients with NMIBC^10,11^. Trained immunity is a form of epigenetic reprogramming of innate immune cells that results from exposure to inflammatory stimuli and is manifested by enhanced inflammatory responses upon re-exposure to homologous or heterologous stimuli^12,13^. Mouse models of bladder cancer have provided evidence that BCG-reprogrammed bone marrow hematopoietic stem and progenitor cells are important in driving innate and adaptive anti-tumour responses^14–16^.

Thus, therapeutic failure of BCG in patients with bladder cancer may be due to defective reprogramming of systemic immunity. Notably, bladder cancer is predominantly a disease of older individuals, with a median age at diagnosis exceeding 70 years^17,18^. This raises the possibility that ageing-associated immune dysfunction fundamentally limits the capacity of patients to mount effective BCG-induced immune responses. Ageing of the immune system is characterized by a chronic low-grade inflammatory state, termed inflammageing, and a progressive decline in immune function linked to immunosenescence^19^. The latter arises from cumulative lifetime exposure to inflammatory, infectious, and genotoxic stressors, resulting in the sustained activation of inflammatory pathways and epigenetic remodelling of hematopoietic stem and progenitor cells^19–24^. A hallmark of immunosenescence and inflammageing is the senescence-associated secretory phenotype, which in myeloid cells and other senescent cells is characterized by the persistent production of pro-inflammatory cytokines in a manner transcriptionally driven by activator protein 1 (AP-1) signalling^25–27^. In myeloid cells, sustained AP-1 activity may therefore reflect a maladaptive immune state in which inflammatory signalling is maintained, thereby limiting the epigenetic plasticity required for effective immune reprogramming^28^.

In the present study, we employed a single-cell multiomics approach to determine whether the transcriptomic and chromatin accessibility profiles of circulating immune cells from patients with high- and intermediate-risk NMIBC provide insight into the therapeutic response to BCG. We found that monocytes from patients who experienced early disease recurrence had evidence of activation of various inflammatory and metabolic pathways linked to ageing-associated immunosenescence, prior to BCG therapy. This immunosenescence signature was accompanied by increased levels of transcripts encoding members of the AP-1 transcription factor complex and increased chromatin accessibility at AP-1 binding sites.

## Results

### Study design and patient stratification of BCG response in NMIBC

To determine whether systemic immune dysfunction characterises BCG failure in patients with NMIBC, we compared the transcriptomic and chromatin accessibility profiles of peripheral blood mononuclear cells (PBMCs) from patients who remained disease-free for at least a year (BCG non-responders; n=8) and patients who suffered disease recurrence within one year of therapy initiation (BCG responders; n=8). Blood samples from these 16 patients were obtained after resection of bladder tumours, immediately before the first instillation of BCG (pre-BCG) and one week after the fifth BCG instillation of induction therapy (post-BCG). Eight patients (six males and two females) suffered recurrence within one year of BCG induction, whereas seven of eight (five males and two females) remained disease-free for at least two years and one male patient was free of disease for over a year but died of a cause unrelated to bladder cancer before the two-year mark. Three patients had stage T1 disease (all male) while 13 had stage Ta disease, 11 patients were high grade and 5 were low grade. Patient characteristics are described in Table 1.

**Table 1.** Patient characteristics.

| Characteristic | BCG Non-Responders<br>n = 8 | BCG Responders*<br>n = 8 |
| --- | --- | --- |
| Age at Consent<sup>1</sup> | 74.0 (9.8) | 73.5 (11.9) |
| Sex |  |  |
| Female | 2 (25%) | 2 (25%) |
| Male | 6 (75%) | 6 (75%) |
| Stage |  |  |
| T1 | 1 (13%) | 2 (25%) |
| Ta | 7 (88%) | 6 (75%) |
| Grade |  |  |
| High with CIS | 0 (0%) | 3 (38%) |
| High | 5 (63%) | 3 (38%) |
| Low | 3 (38%) | 2 (25%) |
| AUA Risk |  |  |
| 2 | 4 (50%) | 3 (38%) |
| 3 | 4 (50%) | 5 (63%) |
| Time to Recurrence (Days)<sup>2</sup> | 135 (73, 305) | NA |

\* Seven of eight BCG responders remained disease-free for more than 24 months. One BCG responder died of a cause unrelated to bladder cancer prior to two years.
<sup>1</sup> Mean (standard deviation)
<sup>2</sup> Median (minimum, maximum)
CIS = carcinoma in situ
AUA Risk = American Urological Association Risk Score

### Single-nuclei multiomic profiling defines the peripheral blood mononuclear cell immune landscape in BCG-treated patients

Single-nuclei multiomic profiling of PBMCs captured both transcriptomic (scRNA-seq) and chromatin accessibility (snATAC-seq) landscapes across approximately 6,500 high quality nuclei per patient per time point (total pre-BCG: 100,523; total post-BCG: 98,129). Joint integration of pre-BCG snRNA-seq and snATAC-seq data using Uniform Manifold Approximation and Projection (UMAP) identified 12 distinct immune cell populations, comprising 43% myeloid and 57% lymphoid cells (Supp. Fig. 1a, Fig. 1a). These populations recapitulated the expected PBMC landscape, including monocyte, dendritic cell, T cell, NK cell, B cell, and plasma cell compartments, and were defined by canonical transcriptional markers (Supp. Fig. 2). Cell identities were further supported by concordant chromatin accessibility profiles, enabling high-confidence annotation across modalities.

**Figure 1.**
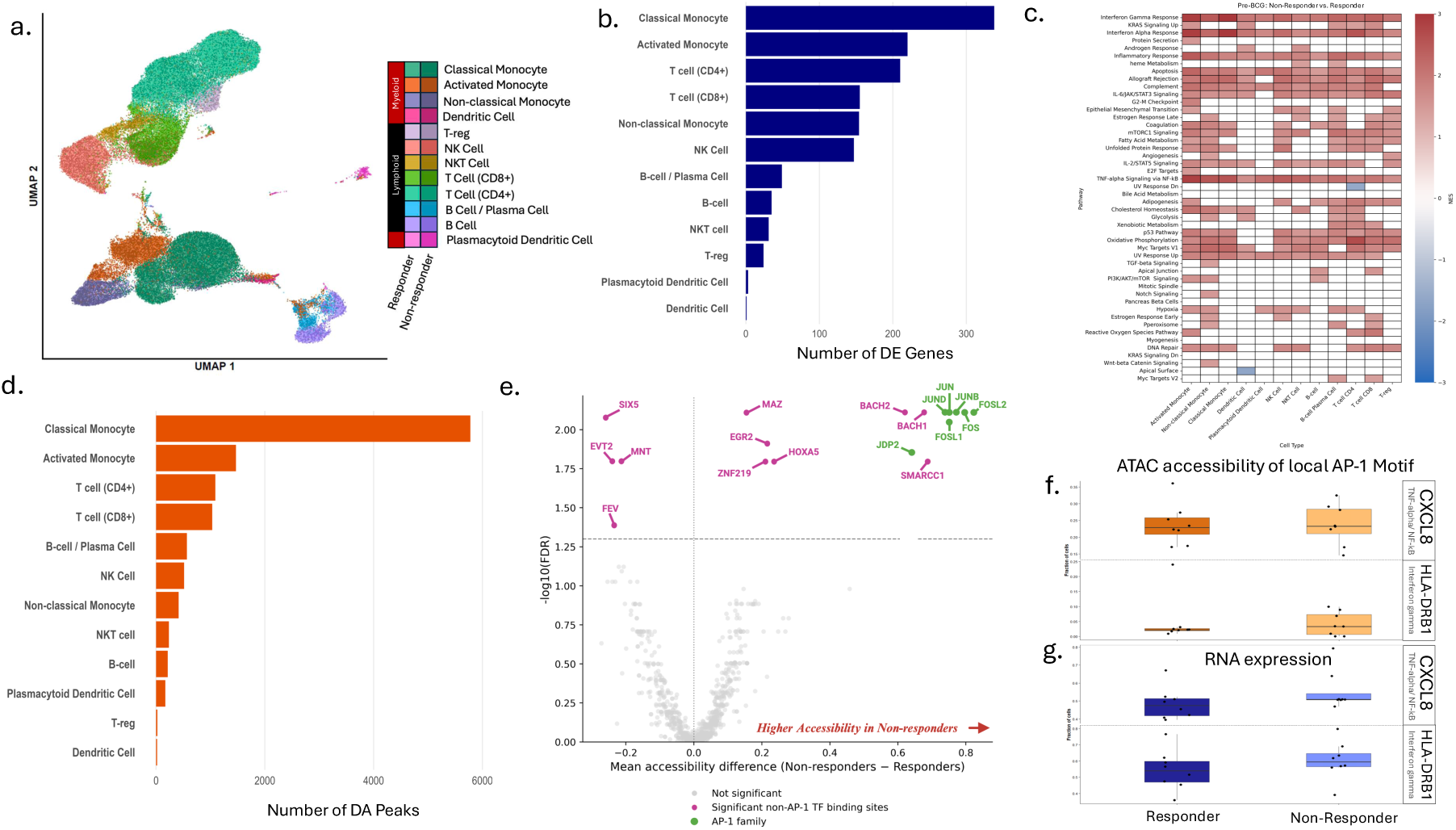
Pre-BCG transcriptional and epigenetic differences in PBMCs identify an AP-1–driven pro-inflammatory chromatin state in patients who fail to respond to therapy. Single-cell multiomic profiling (joint RNA and ATAC) was used to characterize baseline transcriptional and epigenetic differences in PBMCs of patients who subsequently responded to BCG therapy (BCG responders) and those who experienced recurrence (BCG non-responders). (a) UMAP visualization of the joint RNA–ATAC embedding highlights major immune cell lineages, including myeloid and lymphoid compartments, with cellular identities annotated and distributed across both responders and non-responders. (b) Bar plots summarizing the number of differentially expressed (DE) genes (Wilcoxon test on cell-level expression) across cell types between BCG responders and non-responders, prior to BCG therapy. (c) Pseudobulk pathway enrichment analysis (Hallmark gene sets) demonstrates widespread up-regulation of inflammatory and stress-associated pathways across multiple immune cell populations in non-responders compared with BCG responders, including interferon and TNFα/NF-κB signalling, consistent with a baseline pro-inflammatory or immunosenescence phenotype. Bar plots summarizing the number of differentially accessible (DA) chromatin regions across cell types between BCG responders and non-responders, prior to BCG therapy. (e) Differential transcription factor motif accessibility analysis reveals a marked enrichment of AP-1 family motifs in BCG non-responders relative to responders. The x-axis represents the mean accessibility difference (non-responders − responders), while the y-axis denotes statistical significance (−log10 FDR). AP-1 motifs (green) show consistent rightward shifts, indicating increased accessibility in BCG non-responders, whereas non-AP-1 motifs (pink) show more heterogeneous patterns. (f,g) Integration of chromatin accessibility and transcriptional output highlights functional consequences of AP-1 activation at key inflammatory loci. (f) Accessibility at local AP-1 binding sites is increased in non-responders for representative genes associated with enriched pathways in (c), including *CXCL8* (TNFα/NF-κB signalling) and *HLA-DRB1* (interferon-γ response), as measured by the fraction of cells with accessible chromatin. (g) Correspondingly, RNA expression of these genes is elevated in non-responders, linking increased AP-1 motif accessibility to downstream transcriptional activation

**Figure 2.**
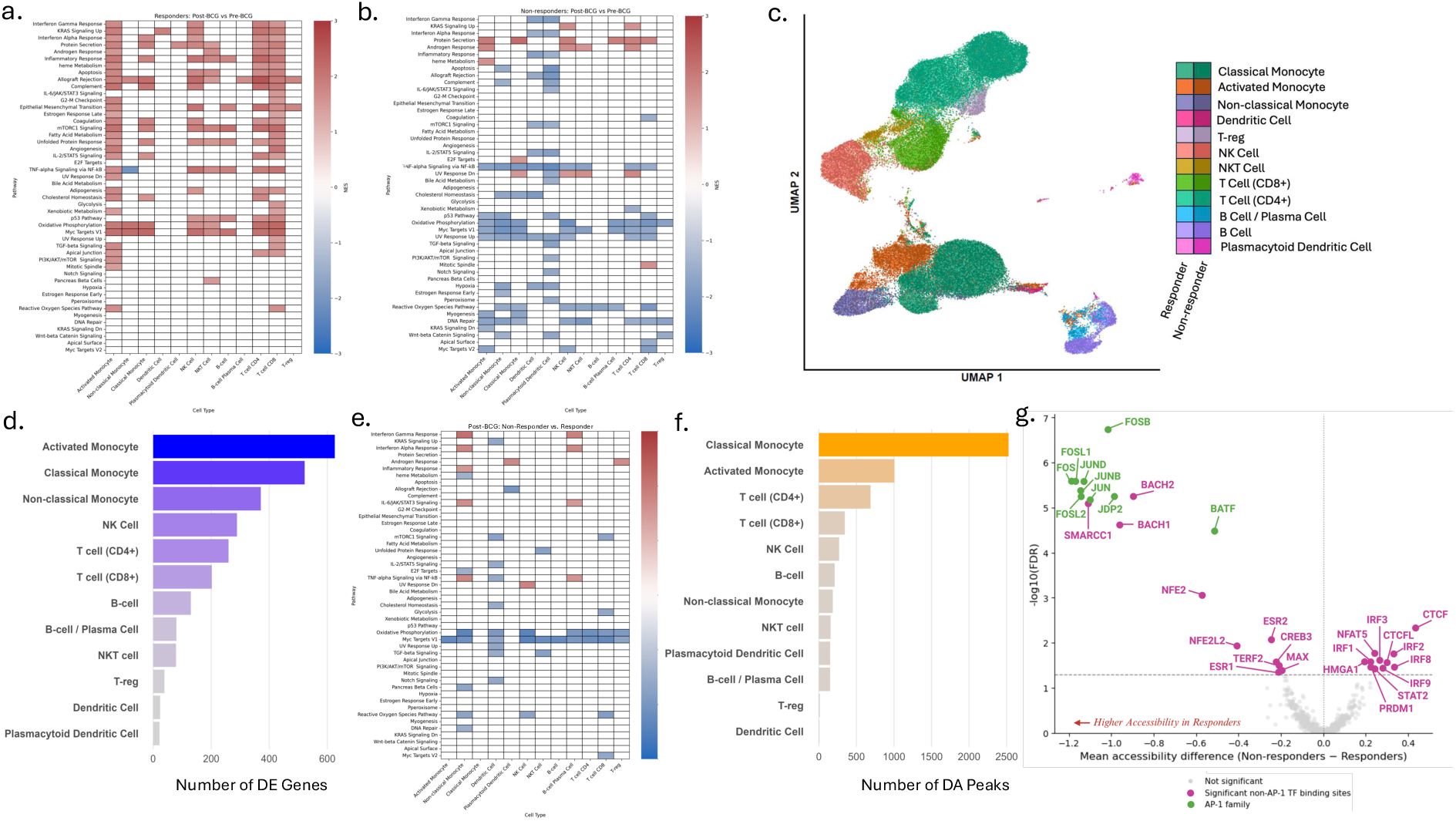
Post-BCG immune remodelling reveals response-associated activation of inflammatory programs and reversal of AP-1–linked chromatin accessibility patterns. (a,b) Longitudinal pathway analysis (post-BCG vs pre-BCG, pseudobulk Hallmark enrichment) reveals divergent immune remodelling between response groups. (a) Responders exhibited coordinated up-regulation of inflammatory, metabolic, and proliferative signalling pathways following therapy. (b) In contrast, non-responders demonstrate widespread down-regulation or failure to induce these same pathways. (c) UMAP visualization of the joint RNA–ATAC embedding highlights major immune cell lineages, including myeloid and lymphoid compartments, with broadly comparable cellular distributions between responders and non-responders following therapy. (d) Bar plots summarizing the number of differentially expressed (DE) genes (Wilcoxon test) on cell-level expression across cell types between responders and non-responders. (e) Heatmap displaying a direct comparison of pseudobulk Hallmark enrichment between non-responders and responders in the post-BCG setting reveals relatively few consistent pathway-level differences between groups. (f) Bar plots summarizing the number of differentially accessible (DA) chromatin regions across cell types between responders and non-responders (g) Differential transcription factor motif accessibility analysis highlights a reversal of pre-treatment regulatory patterns. The x-axis represents mean accessibility difference (non-responders − responders), and the y-axis denotes statistical significance (−log10 FDR). Significant non-AP-1 transcription factor binding sites are shown in pink, AP-1 family motifs in green, and non-significant sites in gray.

### Pre-BCG transcriptional and epigenetic states distinguish BCG responders from non-responders

#### Immunosenescence-associated myeloid transcriptional programs define pre-treatment response stratification

To determine whether differences in response to BCG therapy correlate with pre-existing transcriptional states, an independent joint RNA-ATAC integrated embedding of exclusively pre-BCG PBMC populations was investigated. Notably, BCG responders and BCG non-responders had comparable baseline cellular compositions (all cell types p > 0.5), including similar distributions of lymphoid and myeloid compartments (p = 0.72) before therapy. However, marked differences between these groups were found at the molecular level. Cell-level Wilcoxon-based differential expression analysis comparing responders and non-responders revealed 1,367 genes differing across all assessed cell types, with the bulk of differentially expressed genes (DEGs) detected in the monocyte populations at baseline (classical monocyte DEGs: 338 (∼1%), activated monocyte DEGs: 220 (0.5%), non-classical monocyte DEGs: 154 (< 0.5%)), with the remaining distinctions primarily detected in CD4^+^ T cells (DEGs: 210 (0.5%)), CD8^+^ T cells (DEGs: 155 (<0.5%)), and NK cells (DEGs: 147 (<0.5%)) (Fig. 1b). Across monocyte populations, genes up-regulated in non-responders were enriched for pro-inflammatory and stress-response programs, including cytokines and chemokines (*e.g.*, *CXCL8, CCL2*), NF-κB signalling regulators (*e.g.*, *NFKBIA*, *TNFAIP3*), and AP-1 transcription factors (*e.g.*, *JUNB*, *FOS*, *ATF3*), alongside immediate early response genes associated with cellular stress and immunosenescence (*e.g.*, *IER3*, *DUSP1*, *HSPA6*) (Supp. Fig. 3).

**Figure 3.**
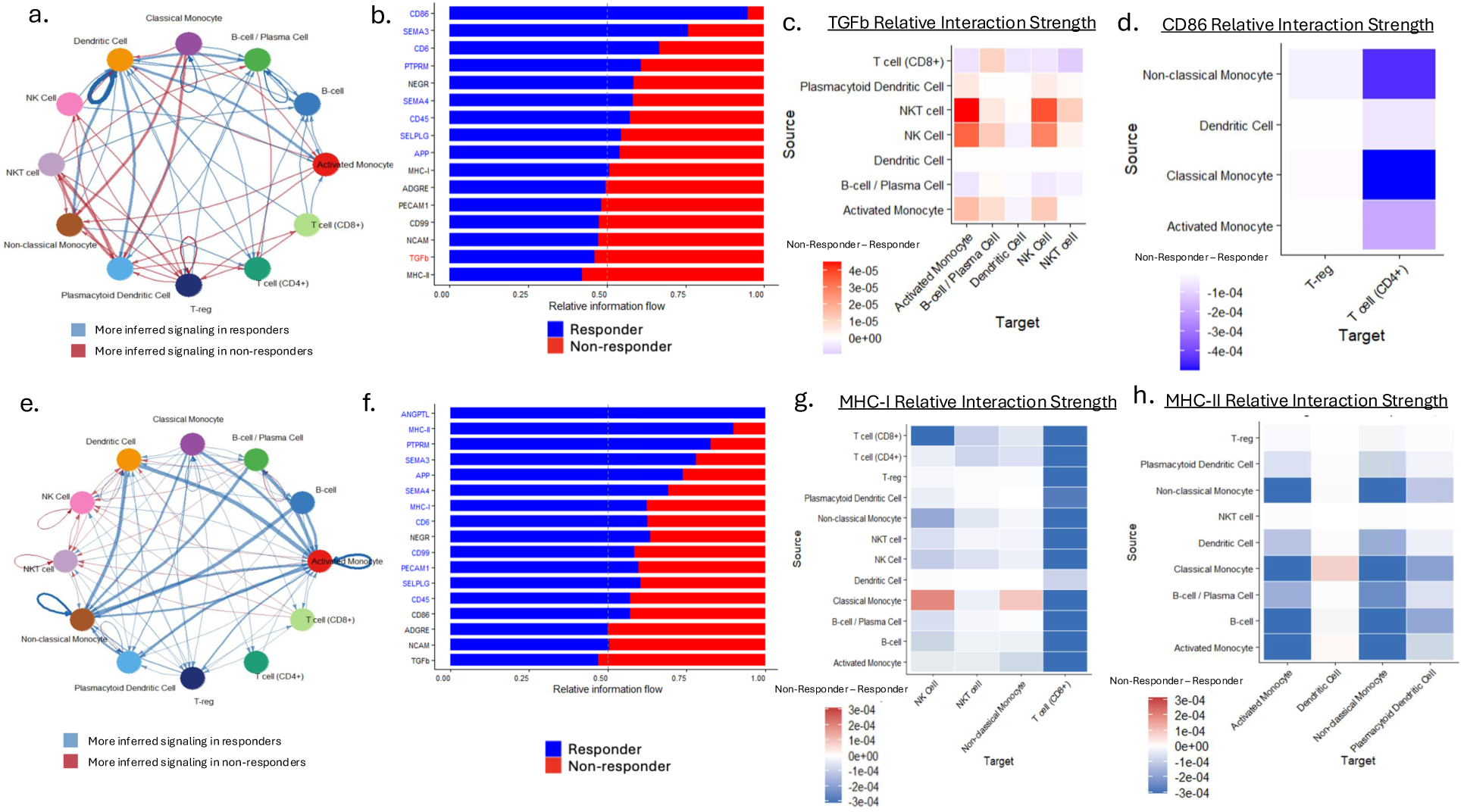
Differential cell–cell communication dynamics before and after BCG therapy reveal opposing baseline immune states and divergent treatment-induced signalling responses. Cell–cell communication analysis was performed using CellChat on PBMC single-cell RNA-seq data, comparing patients who subsequently responded to BCG therapy (BCG responders) versus those who experienced recurrence (BCG non-responders), in both pre- and post-treatment settings. **(a–d) Pre-BCG setting.** (a) Circular network plots depict the differential number of inferred ligand–receptor interactions between BCG responders and non-responders prior to therapy. Node size reflects the number of cells per population, while edge thickness represents the relative number of significant interactions. (b) Differential pathway contribution analysis shows the relative information flow across signalling pathways, partitioned by response group. BCG responders demonstrate increased co-stimulatory signalling, including CD86-associated pathways, whereas BCG non-responders exhibit relatively higher TGFβ signalling, consistent with immunosuppressive and immunosenescence-associated programs. (c,d) Heatmaps illustrate differential cell-type–specific signalling interactions (BCG non-responder minus BCG responder), with blue indicating higher signalling capacity in BCG responders and red indicating higher signalling in BCG non-responders. (c) TGFβ signalling is elevated in non-responders, with broader engagement across innate and lymphoid compartments. (d) In contrast, CD86 signalling is enhanced in BCG responders, particularly involving monocyte-to–T cell interactions. **(e–h) Post-BCG setting.** (e) Circular network plots show an overall increase in the number and strength of inferred interactions in BCG responders following therapy, indicating enhanced intercellular communication in responders. (f) Differential pathway contribution analysis reveals up-regulation of multiple immune activation pathways in BCG responders post-BCG, including MHC-I and MHC-II signalling. (g,h) Heatmaps display differential signalling interactions (non-responder minus responder) for key antigen presentation pathways. (g) MHC-I signalling is broadly increased in BCG responders across multiple sender and receiver populations, including monocyte and dendritic cell compartments. (h) Similarly, MHC-II signalling is enhanced in BCG responders, with strong contributions from classical monocytes and dendritic cells toward lymphoid targets following BCG therapy.

Accordingly, we next assessed whether these transcriptional differences converged at the level of coordinated pathway activity. Consistent with this, pathway analysis of genes up-regulated in BCG non-responders demonstrated enrichment of immune activation and immunosenescence-associated programs. Significant enrichment of interferon gamma and alpha signalling, TNFα signalling via NF-κB, mTORC1 signalling, and MYC target pathways^29^ was observed (Fig. 1c, Supp. Fig. 4). Collectively, the enrichment of interferon, TNFα/NF-κB, and mTORC1 signalling pathways in BCG non-responders is consistent with an immunosenescence-associated inflammatory state characterized by chronic innate immune activation and maladaptive inflammatory remodelling^30,31^.

**Figure 4.**
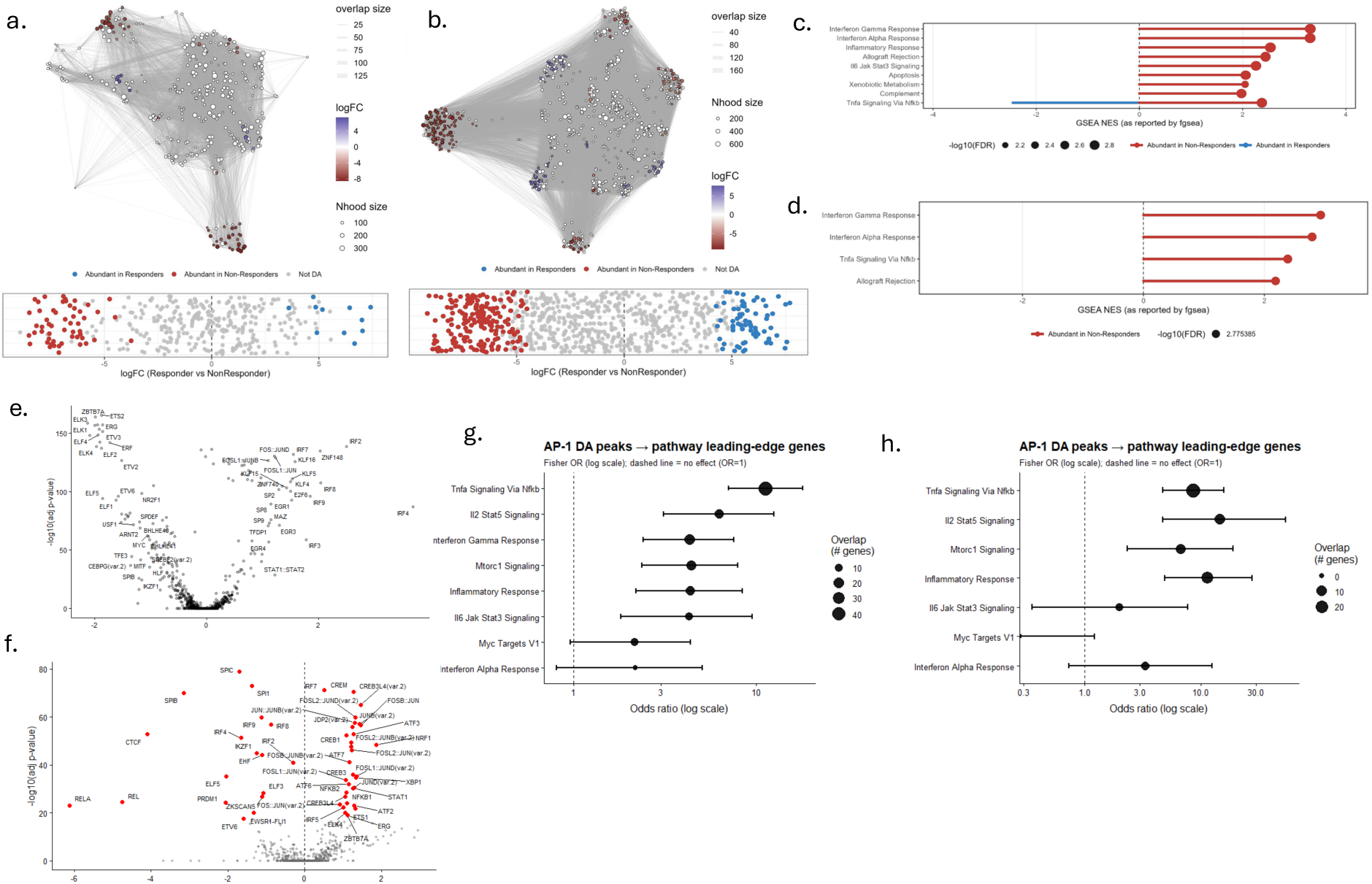
Differential chromatin accessibility and transcriptional programs in monocyte subsets distinguish non-responders from responders in pre-BCG PBMCs. Differential abundance (DA) and regulatory analyses were performed on pre-treatment PBMC single-cell multiomic data to compare patients who subsequently responded to BCG therapy versus those who experienced recurrence. (a,b) Milo-based differential abundance analysis identifies neighbourhoods enriched between groups for activated monocytes (a) and classical monocytes (b). Each node represents a neighbourhood, with colour indicating log fold-change (upregulated in responders; blue vs. upregulated in non-responders; red). Neighbourhoods enriched in non-responders are highlighted, revealing expansion of specific monocyte subpopulations prior to therapy. (c,d) Lollipop plot of Hallmark pathway enrichment demonstrating normalized enrichment scores (NES) and significance (-log10(FDR)) from fgsea analysis comparing the expression of genes upregulated in neighbourhoods that are enriched in (c) activated or (d) classical monocytes of non-responding patients vs. all other cells of that cell type. (e) Volcano plot representing differential accessibility of transcription factor motifs between activated monocyte neighbourhoods up-regulated in non-responding patients vs. all other activated monocytes. (f) Volcano plot representing differential accessibility of transcription factor motifs between classical monocyte neighbourhoods up-regulated in non-responding patients vs. all other classical monocytes. (g,h) Forest plots showing Fisher’s exact test enrichment of AP-1 target genes among the leading-edge genes from the inflammatory pathways defined in (c,d), in (g) activated and (h) classical monocytes derived from cell neighbourhoods up-regulated in BCG non-responding patients. AP-1 target genes were defined as genes located within 500 bp of AP-1–associated differentially accessible peaks. Odds ratios greater than 1 indicate that genes driving pathway enrichment are more likely than expected to be associated with nearby AP-1–linked regulatory regions.

#### Differences in chromatin accessibility reveal AP-1-associated epigenetic priming in BCG non-responders

Consistent with the transcriptional profiles, chromatin accessibility analysis showed a similar pre-BCG pattern, with monocyte populations exhibiting the highest number of differentially accessible regions (DARs). Classical monocytes exhibited the highest number of DARs (n = 5778), followed by activated classical monocytes (n = 1470) and non-classical monocytes (n = 416), whereas substantially fewer differences were observed in lymphoid populations, including CD8^+^ T cells (n = 1033), CD4^+^ T cells (n = 1090), and NK cells (n = 516) (Fig. 1d).

Motif enrichment analysis of these regions identified significant enrichment of transcription factor binding sites associated with the AP-1 family, including *JUN, JUNB, JUND, FOS*, and *FOSL2* (Fig. 1e), which were predominantly associated with regions exhibiting increased accessibility in BCG non-responders. Additional differentially accessible motifs were associated with transcription factors involved in cellular stress responses and lineage-specific regulation, including factors such as *EGR2, MAZ*, and *HOXA5*, although these were less consistently enriched than AP-1–associated motifs. Overall, these differences in chromatin accessibility were most pronounced within monocyte populations pre-BCG and were detectable, albeit to a lesser extent, across other immune cell types. Several of the DARs containing AP-1 motifs were associated with genes that contribute to the inflammatory signalling programs observed in myeloid populations prior to BCG therapy (*e.g., CXCL8* (TNFα via NF-κB) and *HLA-DRB1* (Interferon Gamma Response); local AP-1 motifs DAR adjusted p < 0.005, DEG adjusted p < 0.05), with these differences remaining modest in magnitude but consistently detectable across analyses (Fig. 1f,g).

### Divergent pathway responses to BCG therapy distinguish responders from non-responders

To directly assess BCG treatment–associated changes, pathway analysis was performed using cell-type–specific pseudobulks, comparing patient-matched post- versus pre-BCG samples independently in BCG responders and BCG non-responders. This analysis revealed contrasting patterns of inflammatory, metabolic, and proliferative pathway modulation between the two groups.

In responders, BCG therapy was associated with increased activity of inflammatory pathways, including interferon gamma signalling, complement, IL-2/STAT5 signalling, and TNF-α signalling via NF-κB^29^ (Fig. 2a, Supp. Fig. 5). Metabolic programs, such as oxidative phosphorylation, and proliferative signalling pathways, including mTORC1, and MYC targets, were additionally enriched across myeloid and lymphoid compartments (Fig. 2a, Supp. Fig. 5), consistent with previous reports of systemic immune activation following intradermal BCG vaccination^28^. In contrast, BCG non-responders exhibited a reduction in the activity of the same inflammatory, metabolic, and proliferative pathways following BCG therapy, including decreased interferon gamma response, complement, TNF-α signalling via NF-κB, mTORC1 signalling, and MYC targets^29^ across immune cell populations (Fig. 2b, Supp. Fig. 6).

**Figure 5.**
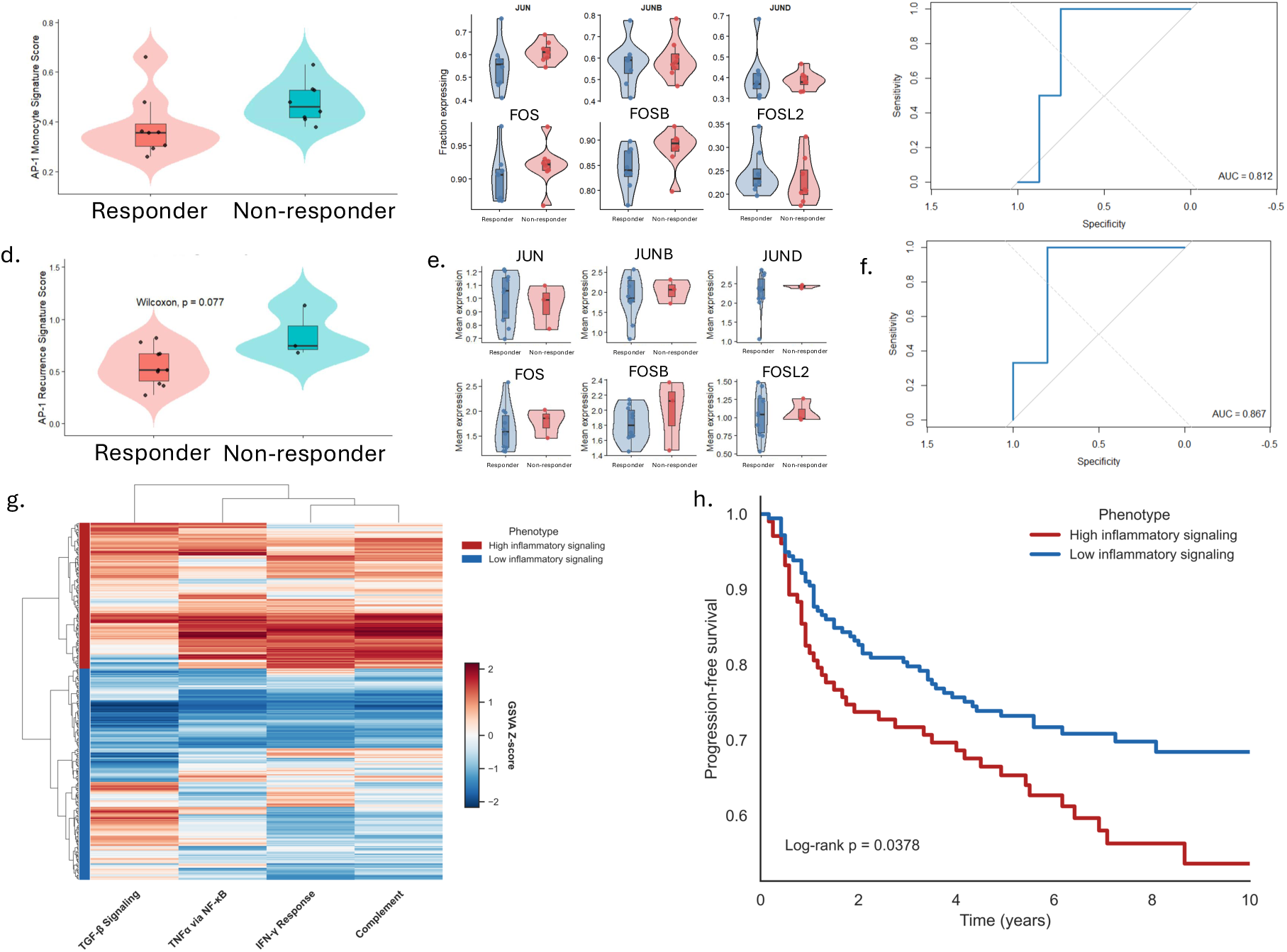
An AP-1–driven immunosenescence signature derived from peripheral myeloid cells predicts BCG response and is associated with poor clinical outcomes in independent bladder cancer cohorts. An AP-1 transcriptional signature was constructed from key AP-1 family transcription factors (JUN, JUNB, JUND, FOS, FOSB, FOSL1, FOSL2) identified as differentially accessible and expressed in pre-BCG myeloid populations (classical and activated monocytes). Signature scores were computed at the single-cell level using Seurat’s module scoring approach, which calculates the averaged, normalized expression of the gene set relative to control gene bins matched for expression. (a,b,c) Validation of the AP-1 signature in PBMC-derived myeloid cells from the study cohort. (a) Violin plots show higher AP-1 module scores in BCG non-responders relative to BCG responders at baseline, consistent with elevated AP-1 activity in patients who fail BCG therapy. (b) Expression of individual AP-1 transcription factors, shown as the fraction of expressing cells, confirms consistent up-regulation of AP-1 components in BCG non-responders, supporting the robustness of the composite signature. (c) Receiver operating characteristic (ROC) analysis demonstrates the ability of this signature to discriminate BCG response status (AUC = 0.812). (d,e,f) Independent validation in a tumour-derived myeloid compartment (macrophages and MDSCs). (d) AP-1 module scores are similarly elevated in non-responders, although the difference does not reach statistical significance, likely reflecting limited sample size. (e) Expression patterns of individual AP-1 transcription factors in tumour-associated myeloid cells recapitulate trends observed in PBMCs, indicating conservation of AP-1–associated programs across systemic and local immune compartments. (f) ROC analysis nonetheless demonstrates strong discriminatory performance (AUC = 0.867). (g,h) Extension of AP-1–associated transcriptional signatures to a large independent bladder cancer cohort (BRS, n=282). (g) Hierarchical clustering of gene set variation analysis scores for pathways identified as associated with the immunosenescence-like phenotype (including interferon-γ response, TNFα/NF-κB signaling, TGF-β Signalling, and Complement Signalling) stratifies patients into distinct high- and low-activity groups. (h) Kaplan–Meier analysis demonstrates significantly worse progression-free survival in patients with high pathway representation, linking AP-1–associated immunosenescence programs to adverse clinical outcomes.

### Distinct BCG-induced remodelling in peripheral immune cells distinguishes BCG responders from BCG non-responders

#### Pro-inflammatory myeloid transcriptional programs distinguish BCG responders from BCG non-responders following BCG therapy

To determine whether differences in response to BCG therapy were maintained following treatment, an independent joint RNA–ATAC integrated embedding of post-BCG PBMC populations was investigated. Notably, BCG responders and BCG non-responders exhibited broadly comparable cellular compositions following therapy (all cell types p > 0.5), including similar distributions of lymphoid and myeloid compartments prior to the final induction dose of BCG (Fig. 2c). In contrast to prior studies of intradermal BCG vaccination conducted in younger individuals^32^, BCG therapy was not associated with evidence of systemic myelopoiesis. Instead, differences between these groups were primarily molecular.

Compared with the pre-BCG setting, transcriptional differences between responders and non-responders following therapy were lower in magnitude. Cell-level Wilcoxon-based differential expression analysis comparing BCG responders and BCG non-responders identified between 350 (∼1%) and 610 (∼1.6%) genes differing across monocyte populations, with the largest number of DEGs detected in activated classical monocytes, followed by classical and non-classical monocytes (Fig. 2d). In contrast, comparatively fewer differences were observed in lymphoid populations, including CD8^+^ T cells, CD4^+^ T cells, and NK cells. Across monocyte populations, genes up-regulated in responders were enriched for pro-inflammatory and immune activation programs, including cytokines and chemokines (*e.g.*, *CCL20*), interferon-stimulated genes (*e.g.*, *IFITM1*), and immune regulatory factors (*e.g.*, *FGL2* and *IL1B*), consistent with transcriptional responses associated with BCG-induced immune activation (Supp. Fig. 7).

Comparative pathway enrichment analysis in the post-BCG setting revealed relatively few pathways that were differentially enriched between the non-responders and responders across immune cell populations. However, MYC signalling^29^ was a notable exception, whereby it was consistently down-regulated in non-responders in both myeloid and lymphoid cell types following BCG administration (Fig. 2e, Supp. Fig. 8).

#### Chromatin accessibility differences following BCG therapy are enriched for AP-1-associated motifs in BCG responders

Consistent with the transcriptional differences observed following BCG therapy, differential chromatin accessibility analysis revealed pronounced epigenetic differences between BCG responders and non-responders, despite similar cellular composition. The greatest number of DARs was detected within monocyte populations, with classical monocytes exhibiting the highest number of DARs (n = 2525), followed by activated monocytes (n = 1006), and substantially fewer differences observed across lymphoid populations (CD8+ T cell; n = 346, CD4+ T cell; n = 688, NK cell; n = 269) (Fig. 2f). These differences were visualized on the joint RNA–ATAC embedding, demonstrating localized regions of increased chromatin accessibility within myeloid compartments in responders.

Motif enrichment analysis of these regions identified significant enrichment of transcription factor binding sites associated with the AP-1 family, including *JUN, JUNB, JUND, FOS, FOSL1*, and *FOSB*, which were predominantly associated with regions exhibiting increased accessibility in BCG responders (Fig. 2g). In contrast to the pre-BCG setting, where AP-1 motifs were enriched in non-responders, these results demonstrate a reversal in the directionality of AP-1–associated chromatin accessibility following therapy. Additional differentially accessible motifs were associated with transcription factors involved in immune activation and interferon signalling, including IRF family members (*e.g.*, *IRF1, IRF3, IRF8*) and STAT family regulators, although these were less consistently enriched than AP-1–associated motifs. These findings parallel the robust transcriptional activation of inflammatory and metabolic pathways observed in BCG responders following BCG administration as well as the down-regulation of similar pathways in BCG non-responders.

### Differential cell-cell communication distinguishes BCG responders from non-responders

To determine whether baseline intercellular signalling differed between patients who subsequently responded to BCG therapy and those who experienced early recurrence, cell-cell communication analysis was performed on pre-BCG PBMCs using CellChat^33^. This analysis identified a greater number of inferred ligand-receptor interactions in BCG non-responders, with increased connectivity involving myeloid populations, particularly activated and classical monocytes (Fig. 3a). Differential pathway contribution analysis, quantified as relative information flow across signalling pathways, demonstrated that this increased signalling in BCG non-responders was distributed across multiple immune-related pathways, rather than reflecting selective enrichment of specific immune activation programs (Fig 3b). Notably, BCG non-responders exhibited modestly increased signalling through MHC-II and TNFβ-associated pathways relative to BCG responders, consistent with reported features of immunosenescence-associated immune activation (Fig. 3b,c). Further, CD86-associated co-stimulatory signalling was reduced in BCG non-responders compared with responders before therapy and was primarily restricted to interactions between myeloid populations and CD4⁺ T cell or regulatory T cell (T-reg) compartments (Fig. 3d).

To determine how these baseline differences in intercellular communication were remodelled following BCG therapy, we next performed CellChat analysis on post-BCG PBMCs. In contrast to the pre-treatment setting, BCG responders exhibited a greater number of inferred ligand–receptor interactions following therapy, with increased network connectivity involving both myeloid and lymphoid populations, particularly classical and activated monocytes (Fig. 3e). Differential pathway contribution analysis demonstrated a shift in inferred signalling activity toward BCG responders, with increased relative information flow across multiple immune-related pathways (Fig. 3f). Notably, BCG responders exhibited enhanced inferred signalling through antigen presentation–associated pathways, including MHC-II and MHC-I, as well as co-stimulatory pathways such as CD86. In contrast, BCG non-responders displayed comparatively reduced inferred signalling across these pathways following therapy. Cell-type–specific analysis of MHC-I signalling interactions further demonstrated increased communication probability in BCG responders, with prominent putative signalling originating from classical and activated monocytes toward multiple immune compartments, including T cell populations (Fig. 3g). In BCG non-responders, MHC-II-associated signalling was comparatively weaker and more restricted across cell types (Fig 3h).

Together, these findings indicate that, whereas BCG non-responders are characterized by a broadly elevated yet diffuse baseline signalling landscape, BCG responders exhibit enhanced and coordinated intercellular communication following BCG therapy, particularly within pathways related to antigen presentation and co-stimulatory signalling.

### AP-1-associated chromatin accessibility within differentially abundant monocyte neighbourhoods characterizes BCG non-responder immune states before BCG therapy

To identify cell-state–specific differences between BCG responders and BCG non-responders, differential abundance (DA) analysis was performed using MiloR^34^ on pre-BCG PBMC multiomic data, independently across all annotated cell types. MiloR defines cell ‘neighbourhoods’ as partially overlapping groups of transcriptionally and epigenetically similar cells within the integrated single-cell embedding^34^. Across the immune compartment, classical and activated monocytes were the only cell populations to exhibit differentially abundant neighbourhoods between BCG responders and BCG non-responders (Fig. 4a,b). Within these monocyte compartments, neighbourhoods enriched in non-responders were identified and further interrogated for their transcriptional and regulatory features. Cells within BCG non-responder–associated neighbourhoods demonstrated increased activity of inflammatory and immunosenescence-associated pathways, including interferon responses and TNFα signalling via NF-κB^29^ (Fig. 4c,d).

To define the regulatory basis of these transcriptional programs, motif enrichment analysis of differentially accessible chromatin regions within these neighbourhoods was performed. This analysis revealed significant enrichment of transcription factor binding sites associated with the AP-1 family, including FOS-, JUN-, and ATF-family members, in both activated and classical monocytes (Fig. 4e,f). These AP-1–associated regulatory elements were preferentially enriched within non-responder–associated neighbourhoods compared to other monocyte populations. Integration of chromatin accessibility with pathway-level transcriptional outputs further revealed an enrichment of AP-1 differentially accessible peaks within the promoter and enhancer regions of leading-edge genes in differential pathways, including interferon gamma, TNFα signalling via NF-κB and inflammatory response^29^ within activated and classical monocyte subpopulations that are enriched in non-responders, as quantified by Fisher’s exact test (Fig 4. e-h). Together, these findings demonstrate that classical and activated monocytes represent the primary cell populations exhibiting differential abundance between responders and non-responders prior to BCG therapy, with non-responder-associated neighbourhoods characterized by AP-1-linked chromatin accessibility and enrichment of inflammatory and interferon signalling programs.

### AP-1–associated myeloid regulatory programs and inflammatory signalling define response and survival outcomes across independent bladder cancer cohorts

#### A single-cell–derived AP-1–associated myeloid gene signature stratifies BCG response

An AP-1 myeloid gene signature comprising *JUN, JUNB, JUND, FOS, FOSB* and *FOSL2* was identified from genes linked to differentially accessible AP-1-enriched regulatory regions and up-regulated in BCG non-responders (Fig. 5a,b). Single-cell module scoring revealed that this AP-1 signature was significantly elevated in patients who subsequently did not respond to BCG therapy compared to those who did (Wilcoxon p=0.03792; Fig. 5a). At the patient level, aggregation of signature scores demonstrated clear separation between BCG responders and non-responders, with consistently higher scores observed in recurrence cases. Consistent with this separation, receiver operator characteristic (ROC) analysis demonstrated strong discriminatory performance of the AP-1 signature for predicting BCG response, with an area under of the curve (AUC) of 0.812 (Fig. 5c).

The observed enrichment of AP-1 activity was validated by assessing the coordinated expression of its core complex components (e.g. *JUN, JUNB, JUND, FOS, FOSB, FOSL2*) in an independent cohort^35^ with available pre-BCG tumour-derived single-cell RNA sequencing data (Fig 5d,e). We specifically focused on macrophage populations from patients who subsequently responded to BCG therapy (n = 10) and those who experienced recurrence (n = 3) within 1 year. Transcripts of the AP-1 complex components were elevated in macrophages from BCG non-responders compared to responders, although this difference did not reach statistical significance (Wilcoxon p-value = 0.077; Fig. 5d). Similarly, at the patient level, ROC analysis was performed on the aggregated expression of the AP-1 complex transcripts to assess separation between the outcome groups. This yielded an AUC of 0.867 (Fig. 5f) to suggest AP-1 complex expression retained discriminatory capacity in the independent cohort.

#### AP-1-associated inflammatory signalling programs define patient subgroups with distinct progression-free survival

To assess whether the immunosenescence-associated signalling programs identified in circulating myeloid populations are recapitulated within the pre-treatment local tumour environment, we evaluated pathway-level activity in bulk RNA-sequencing data from the BRS cohort (n =282)^36^. Given that the AP-1–associated transcriptional signature derived from our single-cell analyses is largely restricted to monocyte and myeloid compartments, direct projection of this signature into bulk tumour data is inherently limited by cellular heterogeneity. Instead, we focused on downstream inflammatory pathways enriched in association with this phenotype, specifically the hallmark interferon gamma response, TNFα signalling via NF-κB, complement, and TGF-β gene sets. Gene set variation analysis (GSVA)^37^ was applied to quantify the relative enrichment of these pathways across tumours, enabling sample-wise estimation of pathway activity independent of absolute expression levels.

Unsupervised hierarchical clustering of GSVA scores using Ward’s linkage identified two dominant phenotypic groups characterized by concordant high or low activity across all three pathways, hereafter referred to as “high inflammatory signalling” and “low inflammatory signalling” tumours (Fig. 5g). Stratification of patients based on these clusters revealed a significant difference in progression-free survival, with high inflammatory signalling tumours associated with inferior outcomes (log-rank p = 0.0378) (Fig. 5h). These findings indicate that, despite the cell type–specific nature of the underlying transcriptional program, the broader inflammatory signalling milieu linked to AP-1–driven immunosenescence is preserved at the tumour level and captures clinically relevant variation in disease progression. Collectively, this supports the notion that heightened baseline inflammatory signalling, potentially reflective of a chronically activated or immunologically constrained state, may contribute to impaired anti-tumour immunity and adverse clinical trajectories in NMIBC.

Together, these findings support the reproducibility of the AP-1-associated myeloid transcriptional program across compartments and indicate that this signature captures biologically consistent differences associated with BCG response in both peripheral and tumour-resident myeloid cells prior to therapy.

## Discussion

Here we provide evidence that patients who experience early recurrence of NMIBC exhibit an immunosenescence-like phenotype prior to BCG induction immunotherapy. These patients harbour distinct populations of circulating monocytes with transcriptomic and chromatin accessibility profiles that are consistent with a state of chronic activation of inflammatory and metabolic pathways. This immunosenescence-like phenotype is indicative of a pre-existing, systemically dysregulated immune state.

Our study identifies increased expression of AP-1 transcription factors and the enrichment of AP-1 binding site accessibility in myeloid cells from patients who experienced early disease recurrence following BCG therapy. Given the established role of AP-1 in chronic inflammatory signalling and immune ageing^27^, these findings indicate that heightened AP-1 activity is closely linked to the immunosenescence-like phenotype that characterizes pre-BCG non-responders. Indeed, AP-1 is known as a ‘pioneer’ transcription factor that drives acquisition of oncogene-induced senescence and the senescence-associated secretory phenotype^27^. The chronic AP-1 activation we observed in BCG non-responders may therefore reinforce persistent inflammatory programs that limit responsiveness to BCG immunotherapy^38,39^. Notably, persistent inflammatory activation has been linked to impaired immunotherapeutic responses through the induction of T-cell dysfunction and exhaustion^39^.

Activation of Jun/Fos AP-1 family members is a conserved feature of immune ageing associated with chronic inflammation and functional remodelling of monocytes and macrophages^25^. Our findings define a transcriptional signature comprising core components of the JUN/FOS complex (*i.e.*, *JUN*, *JUNB*, *JUND*, *FOS*, *FOSB* and *FOSL2*) that demonstrates robust discriminatory capacity for predicting BCG response. This signal was reproducible in an independent tumour-derived cohort^35^ whereby coordinated expression of AP-1 components in macrophage populations retained strong classification performance despite limited sample size. Given the cell type-restricted nature of the program, validation in the BRS cohort^36^ was performed at the pathway level which revealed that patients with tumours characterized by immunosenescence-associated high interferon and TNFα, complement, and TGF-β signalling had significantly poorer progression-free survival. These findings are consistent with prior studies implicating sustained AP-1 activity in chronic inflammatory signalling and dysfunctional myeloid states^25^.

In contrast to the elevated AP-1 expression and accessibility observed in BCG non-responders prior to treatment, BCG responders exhibited increased AP-1 expression and chromatin accessibility following therapy. This finding is suggestive of a distinct role for BCG-induced AP-1 activation during productive anti-tumour responses. In support of this, AP-1 has been shown to be necessary for BCG-induced, TNFα-driven, inflammatory programs in macrophages and monocytes^40–42^. Our findings show that BCG responders retain the capacity to up-regulate these AP-1-associated inflammatory programs in myeloid cells following therapy. Conversely, non-responders appeared to enter treatment with a chronically elevated AP-1-associated inflammatory state that failed to undergo further transcriptional and epigenetic remodelling following BCG exposure, instead demonstrating attenuation of inflammatory and metabolic signalling after therapy. This reversal in AP-1-associated chromatin accessibility may therefore reflect preserved immune plasticity in responders versus a state of inflammageing-associated “immune paralysis”^43^ in non-responders, whereby chronic baseline AP-1 activation is uncoupled from therapeutically effective immune reprogramming.

Our analysis of intercellular communication further supports a model in which BCG responsiveness is determined primarily by its coordinated ability to activate key inflammatory pathways. Prior to therapy, BCG non-responders exhibited a broadly elevated, yet diffuse signalling landscape characterized by heightened myeloid ligand expression and modest enrichment of pathways associated with antigen presentation and inflammatory signalling. This pattern is consistent with an immunosenescence-like state of chronic immune activation that is maintained without effective coordination^19,44^. The ineffective coordination therefore may constrain downstream therapeutic response. In contrast, following BCG therapy, BCG responders exhibited a marked reorganization of intercellular communication networks with increased connectivity across myeloid and lymphoid compartments. This heightened intercellular communication was associated with coordinated up-regulation of antigen presentation pathways (*e.g.*, MHC-I and MHC-II) alongside enhanced co-stimulatory signalling through CD86. Notably, these signalling interactions not only increased in magnitude but were structured across defined ligand-receptor axes, particularly those involving classical and activated monocytes and T cell populations. This inferred interaction is consistent with effective antigen presentation and adaptive immune priming. In this context, the elevated signalling observed in non-responders before BCG therapy likely reflects a maladaptive, functionally disordered, immune state that lacks the epigenetic and transcriptional plasticity required for effective reprogramming following therapy.

The identification of differentially abundant monocyte neighbourhoods in BCG non-responders further refines this model by implicating discrete myeloid cell states, rather than global immune composition, as key determinants of therapeutic outcome. Consistent with a prior report demonstrating that circulating monocyte subsets are major mediators of systemic inflammatory amplitude and predictors of immunotherapeutic response^15^, differential abundance was restricted to classical and activated monocytes. The differentially abundant neighbourhoods associated with a lack of response to BCG therapy were characterized by enrichment of interferon and TNF-alpha/NFκB signalling programs alongside AP-1-associated chromatin accessibility. This suggests that the expansion of transcriptionally primed myeloid subpopulations in the bone marrow drives the development of the immunosenescence phenotype. Notably, AP-1 motif enrichment was localized to regulatory elements of leading-edge genes in these inflammatory pathways, supporting a direct coupling of AP-1-driven chromatin landscape and associated transcriptional output. This finding substantiates the role of AP-1 as a context dependent regulator of enhancer activity in myeloid cells, specifically under conditions of chronic stimulation whereby AP-1 contributes to the stabilization of inflammatory gene expression programs and altered cellular responsiveness^25^.

Although our integrative single-nuclei transcriptomic and chromatin accessibility analyses provide novel insights into systemic immune states associated with BCG failure, some limitations merit consideration. First, the relatively small cohort limits statistical power and may constrain the generalizability of our findings, particularly given the known heterogeneity of immune responses in NMIBC^45^. Further, validation of the AP-1-driven immunosenescence signature was performed in a similarly small independent cohort of tumours from patients treated with BCG. Scarcity of single cell sequencing profiles from PBMCs in the context of BCG immunotherapy emphasizes the need for further validation. While our findings implicate immunosenescence-associated transcriptional programs in mediating BCG response, functional validation of this relationship requires experimental modelling. The implementation of mouse models that recapitulate age-associated immune dysfunction will enable testing of interventions aimed at targeting immunosenescence.

The AP-1-associated signature identified here could serve as a minimally invasive biomarker to stratify patients and inform therapy. For example, our findings support the exploration of alternative clinically approved therapeutic strategies, such as gemcitabine/docetaxel-based regimens^46^, for patients identified as unlikely to benefit from BCG therapy. Furthermore, since immunosenescence can be targeted with senolytics or senomorphics, such as rapamycin, our study provides a rationale for clinical studies to determine whether these drugs can improve BCG responses in patients who would otherwise fail BCG therapy. Indeed, daily rapamycin was found to enhance BCG-specific immune responses and to be well tolerated and safe in patients with NMIBC treated with BCG^47^. The pre-clinical discovery showing that senolytic therapy enhances the efficacy of Programmed Death Ligand 1 checkpoint inhibition in a mouse MC38 colorectal cancer model^38^ also supports its potential to improve the efficacy of emergent therapeutic approaches, such as pembrolizumab, for the treatment of NMIBC^48^. Lastly, the relevance of our findings may extend beyond NMIBC to other cancer types by providing insight into why many elderly patients with cancer often fail to respond effectively to immunotherapy. Leveraging this foundational concept could ultimately inform strategies to improve treatment response and outcomes for older adults with cancer.

## Materials and Methods

### Study participants

For transcriptomic and genome-wide chromatin accessibility analysis at the single nuclei level, 16 patients with intermediate or high-risk NMIBC were recruited prior to initiation of BCG induction therapy and after informed consent and approval by the Queen’s University Health Sciences and Affiliated Hospitals Research Ethics Board (approval number UROL-282-13). Eight patients suffered disease recurrence within a year of BCG therapy initiation (BCG non-responders) and eight patients remained disease free for at least two years (BCG responders). Guidelines from the American Urological Association informed the risk stratification, surveillance and adjuvant therapies, and definitions of recurrence in our cohort^49^. Exclusion criteria included exposure to BCG within the previous three years, as such exposure reduces the likelihood of a response to BCG and complicates the assessment of immunological responses in the BCG naïve setting^50^. Patient characteristics are described in Table 1. Patients with high-grade T1 disease were re-resected to ensure accurate staging.

### Preparation of single-nuclei multiome libraries

Prior to nuclei isolation, one vial of PBMCs per participant was thawed and processed according to 10x Genomics’ demonstrated protocol for PBMC nuclei isolation. After thawing, PBMCs were transferred to a 50-mL conical tube and sequentially diluted by incremental 1:1 volume additions of medium (10% FBS in RPMI 1640) for a total of five times at room temperature. After equilibration in separation buffer (PBS + 1% BSA + 2 mM EDTA), cells were resuspended in 100 μL separation buffer 40 μL each of Erythrocyte Depletion Beads and Granulocyte Depletion Microbeads (Miltenyi Biotec) were added to the cell suspension and incubated at room temperature for 5 minutes. Volumes were then completed to 2 mL with separation buffer and passed through LS columns on a QuadroMACS separator (Miltenyi Biotec). The flow-through (purified PBMCs) was collected and used for nuclei isolation.

After equilibration in PBS + 0.04% BSA, subsequent steps were performed at 4℃. One million cells were pelleted at 300g for 5 minutes and resuspended in 100 μL ice-cold lysis buffer (10 mM Tris-HCl pH 7.4, 10 mM NaCl, 3 mM MgCl2, 0.1% Tween-20, 0.1% IGEPAL, 0.01% digitonin, 1% BSA, 1 mM DTT, 1 U/μL RNase inhibitor). After 3 minutes of lysis, one mL ice-cold wash buffer (10 mM Tris-HCl pH 7.4, 10 mM NaCl, 3 mM MgCl2, 0.1% Tween-20, 1% BSA, 1 mM DTT, 1 U/μL RNase inhibitor) was added and nuclei were pelleted at 500g for 5 minutes. After resuspension for the second wash (1 mL wash buffer each time), nuclei were passed through a 40 μm cell strainer (VWR). Nuclei were resuspended in 50 μL and counted using a Countess 3 cell counter (Invitrogen). Transposition, GEM generation and barcoding, and library preparations were performed according to 10x Genomics Chromium Next GEM Single Cell Multiome ATAC + Gene Expression protocol using a targeted recovery of 10,000 nuclei.

Libraries were sequenced on an Illumina NovaSeq 6000 instrument at McGill Genome Centre (Montréal, QC, Canada) or Illumina NovaSeq X instrument at Princess Margaret Genomics Centre (Toronto, ON, Canada). Resulting FASTQ files were aligned to the GRCh38 (hg38) reference genome and quantified using Cell Ranger ARC (v2.0.2) count to generate BAM files and matrices used in subsequent analyses.

### Single-nuclei transcriptomics analysis

#### Preprocessing and Quality Control

Single-nuclei RNA sequencing data were processed using Scanpy (v1.11.5)^51^. Gene-barcode matrices generated by Cell Ranger Arc (v2.0.2) were imported into AnnData objects, and samples were combined for downstream analysis. Quality control was performed on a per-sample basis to account for variability in sequencing depth and technical composition. Nuclei were filtered based on gene complexity (minimum 300 detected genes), total unique molecular identifier (UMI) counts, mitochondrial transcript proportion, and ribosomal transcript proportion. Thresholds for mitochondrial (18–35%) and ribosomal (3–8%) content were determined empirically for each sample (Supp. Fig. 11). Mitochondrial transcripts and genes detected in fewer than 150 nuclei were excluded. Putative doublets were identified using Scrublet (v0.2.4)^52^ and removed before downstream analysis (Supp Fig. 8b). Counts were library-size normalized (counts per million; CPM) and log transformed. Highly variable genes (n = 2500) were identified using a batch-aware variance modelling approach, in which gene-wise variability was estimated within each sample and aggregated to minimize batch-driven feature selection. Principal component analysis (PCA) was performed on the scaled data, retaining the top 50 principal components for downstream analysis. To account for inter-sample batch effects, the data were integrated using Harmony (v0.0.9)^53^.

#### Dimensional Reduction, Clustering, and Cell Annotation

A k-nearest neighbour (kNN) graph was constructed on the integrated low-dimensional embedding and visualized on a uniform manifold approximation and projection (UMAP) plot using Scanpy (v1.11.5)^51^. Clustering was performed using the Leiden algorithm^54^ across a range of resolutions (0.02–2.0) to assess cluster stability and hierarchical structure. A representative resolution was selected for downstream analyses based on cluster separation and biological interpretability (Supp. Fig. 2). Cell clusters were annotated based on canonical marker gene expression and cluster-specific differential expression patterns, informed by established lineage-defining markers (Supp. Fig. 2).

#### Wilcoxon-Based Cell-level Differential Expression

Cell-level differential expression analysis was additionally performed using the Wilcoxon rank-sum test implemented in Seurat (v5.4.0)^55^ to identify transcriptional differences between groups at single-cell resolution. Differential expression was conducted independently within each annotated cell type across four comparisons, including cross-sectional comparisons between responders and non-responders at both pre- and post-BCG timepoints, as well as longitudinal comparisons of post- versus pre-BCG samples within each response group. Gene expression values were log-normalized prior to testing, and differential expression statistics were calculated on a per-cell basis using Seurat’s FindMarkers function^55^. P-values were adjusted for multiple testing using the Benjamini–Hochberg procedure, and resulting statistics, including average log2 fold changes and adjusted p-values, were used for downstream analyses.

#### Pseudobulk Differential Expression and Pathway Enrichment Analysis

To enable robust differential expression analysis while accounting for patient-level variability, raw single-nuclei transcript counts were aggregated into pseudobulk profiles by summing gene expression across cells of the same type within each sample, and these pseudobulk counts were leveraged for downstream analyses.

Differential expression analysis was performed using DESeq2 (v1.50.2)^56^ independently within each cell type across four comparisons. Cross-sectional differences between responders and non-responders were assessed at both pre- and post-BCG timepoints using an unpaired design. Longitudinal changes associated with BCG therapy were evaluated within each outcome group using a paired design, comparing post- versus pre-treatment samples while controlling for inter-individual variability. Differential expression was assessed using the Wald test, and p-values were adjusted for multiple testing using the Benjamini–Hochberg procedure. Resulting statistics, including log2 fold changes and test statistics, were used for downstream analyses.

Pathway enrichment analysis was performed using a pre-ranked gene set enrichment approach implemented in Python package, gseapy (v1.1.11)^57^. Genes were ranked based on signed statistics derived from differential expression results for each comparison. Enrichment was assessed against the ‘hallmark gene sets’ from the Molecular Signatures Database (v2026.1.Hs)^29^, and analysis was conducted independently for each of the four comparisons. Normalized enrichment scores (NES) and adjusted p-values were used to determine pathway significance.

### Single-nuclei chromatin accessibility analysis

#### Preprocessing and Quality Control

Single-nuclei ATAC-seq data were processed using ArchR (v1.0.2)^58^. Fragment files generated by Cell Ranger ARC (v2.0.2) were used to construct Arrow files for each sample. During Arrow file generation, nuclei were filtered to retain those with transcription start site (TSS) enrichment ≥ 6 and ≥ 5,000 unique fragments. Genome-wide accessibility was quantified using a tile matrix, and gene activity scores were computed using a gene score matrix.

Quality control metrics, including transcription start site (TSS) enrichment, fragment counts, nucleosomal signal, and fragment size distributions, were evaluated across samples to assess chromatin accessibility quality and sequencing complexity (Supp. Fig. 12c,f). Nucleosomal signal was calculated as the ratio of mononucleosomal to nucleosome-free fragments for each nucleus and was used to identify low-quality nuclei with poor chromatin accessibility profiles. Nuclei with nucleosomal signal values greater than 2 were excluded from downstream analysis (Supp. Fig. 12d). Doublets were identified using ArchR’s simulation-based doublet detection approach^58^ and removed prior to downstream analysis, resulting in the exclusion of 32,279 nuclei (11.3%) across all samples (Supp. Fig. 12b).

#### Dimensionality Reduction, Batch Corrections and Clustering

Dimensionality reduction was performed using iterative latent semantic indexing (LSI) on the tile matrix to capture major sources of variation in chromatin accessibility. To account for inter-sample batch effects, the low-dimensional representation was integrated using Harmony^53^. A k-nearest neighbour (kNN) graph was constructed based on the corrected embedding, and cells were visualized using uniform manifold approximation and projection (UMAP).

Clustering was performed using the Leiden algorithm^54^ at a resolution of 0.15 to define broad chromatin accessibility–based cell populations. To enable consistent cell type annotation across modalities, cluster identities were aligned to matched single-nuclei transcriptomic profiles by barcode correspondence, and cell labels were assigned accordingly.

#### Peak Calling and Transcription Factor Deviation Analysis

Accessible chromatin regions were identified using external peak calling with MACS2 (v2.2.9.1)^59^ on pseudobulked fragment profiles generated for each cell type and sample. Resulting peak sets were imported into the ArchR project and unified to generate a non-overlapping peak set, from which a peak-by-cell accessibility matrix was constructed for downstream analyses.

Transcription factor motif accessibility was quantified using chromVAR-based (v1.12.0)^60^ deviation analysis implemented in ArchR^58^. Motif annotations, from JASPAR2020 (v0.99.10)^61^, were incorporated using curated position weight matrices, and deviations in accessibility across peaks containing shared motifs were computed per cell while correcting for technical biases, including GC content and sequencing depth. Differential transcription factor deviation analysis was performed independently within each cell type across four comparisons: (i) responders versus non-responders prior to BCG therapy, (ii) responders versus non-responders following BCG therapy, (iii) post-versus pre-treatment within responders, and (iv) post- versus pre-treatment within non-responders. Statistical significance was assessed using group-wise testing with multiple testing correction, and transcription factors with significant deviation differences were retained for downstream interpretation.

### Single-nuclei multiomics integration analysis

#### Joint RNA-ATAC preprocessing and Integration

Single-cell multiomic (RNA and ATAC) embeddings were derived from the independently processed transcriptomic and chromatin accessibility analyses described above using Signac (v1.17.1)^62^. Briefly, principal component analysis (PCA) of gene expression data and latent semantic indexing (LSI) of chromatin accessibility data were used as modality-specific low-dimensional representations. These embeddings were integrated using a weighted nearest neighbour (WNN) approach implemented in Seurat^55^, generating a joint representation of cellular state that captures complementary transcriptional and epigenetic information. The resulting multimodal embedding was used for downstream visualization and neighbourhood-based analyses, with cells projected into two-dimensional space using UMAP.

#### Neighbourhood-Based Differential Abundance Analysis

Neighbourhood-based differential abundance analysis was performed using MiloR (v2.6.0)^34^ to identify localized changes in cellular composition associated with BCG response. To minimize confounding by global shifts in cell-type abundance, analyses were performed separately for each major cell type (n = 13). A kNN (k=30) graph was constructed from a subset of the integrated multimodal joint embedding. Cellular neighbourhoods were defined using MiloR^34^ by sampling 5% of cells as index nodes and aggregating their local graph neighbourhoods. These neighbourhoods represent partially overlapping groups of transcriptionally and epigenetically similar cells, with neighbourhood size and composition determined by the density of cells within the integrated embedding.

Differential abundance testing was conducted independently for each timepoint (pre-BCG and post-BCG), comparing responders and non-responders. A generalized linear model framework implemented in MiloR^34^ was used to model neighbourhood cell counts while accounting for inter-sample variability. P-values were adjusted using spatial false discovery rate (SpatialFDR) to account for multiple testing across overlapping neighbourhoods, and neighbourhoods with SpatialFDR < 0.05 were considered significantly differentially abundant. Significant neighbourhoods were further stratified based on the direction of enrichment (log fold-change), enabling identification of neighbourhoods expanded in non-responders versus responders and vice versa.

#### Functional and Regulatory Characterization of Differential Neighbourhoods

Differential neighbourhoods were functionally characterized by integrating transcriptional and chromatin accessibility features. Pathway activity was quantified using gene set scoring in Seurat (v5.4.0)^55^ with ‘Hallmark gene sets’ from the Molecular Signatures Database (v2026.1.Hs)^29^. Per-cell module scores were computed and subsequently averaged across cells within each neighbourhood to generate neighbourhood-level pathway activity profiles. Transcription factor activity was quantified using chromVAR (v1.12.0)^60^ deviation analysis, and differential motif accessibility was assessed between neighbourhood-defined populations and background cells using non-parametric testing.

To define the regulatory basis of these transcriptional programs, chromatin accessibility profiles were interrogated within differential neighbourhoods using the ATAC assay within the integrated Seurat/Signac (v1.17.1)^62^ framework. Differential chromatin accessibility analysis was performed at the peak level by comparing accessibility between cells within non-responder–associated neighbourhoods and background cells from the same cell type using FindMarkers with logistic regression, accounting for sequencing depth and technical covariates. Peaks were annotated to their nearest genes to facilitate integration with transcriptional outputs.

Transcription factor activity was inferred using motif enrichment analysis implemented in Signac^62^, leveraging position weight matrices from the JASPAR2020 database (v0.99.10)^61^. Motif enrichment within differentially accessible peaks was assessed using hypergeometric testing to identify transcription factor binding sites overrepresented in non-responder–associated neighbourhoods. This analysis identified significant enrichment of AP-1 family motifs, including members of the FOS, JUN, and ATF families, within both classical and activated monocyte neighbourhoods enriched in non-responders.

To further link regulatory elements with observed transcriptional programs, peaks containing AP-1–associated motifs were intersected with promoter and enhancer regions of genes contributing to leading-edge subsets of enriched pathways identified through gene set scoring. Enrichment of AP-1–associated peaks within promoter and enhancer regions of leading-edge genes from differentially enriched pathways was quantified using Fisher’s exact test implemented in the stats package in R (v4.4.1).

#### AP-1 Monocyte Signature Scoring and Patient-Level Analysis

To quantify AP-1–associated transcriptional activity within circulating myeloid populations prior to BCG therapy, a targeted gene signature was constructed using canonical AP-1 transcription factors (JUN, JUNB, JUND, FOS, FOSB, FOSL2). Analysis was restricted to classical and activated monocytes derived from pre-BCG single-nuclei transcriptomic data. Per-cell module scores were computed using the AddModuleScore function implemented in Seurat^55^, which calculates the average expression of the gene set of interest subtracted by aggregated expression of control gene bins matched for expression levels.

To account for inter-individual variability and enable patient-level comparisons, module scores were aggregated by computing the mean AP-1 signature score across all monocytes within each sample. Resulting patient-level scores were grouped according to clinical outcome (BCG responders versus non-responders). Differences in signature activity between outcome groups were assessed using a two-sided Wilcoxon rank-sum test.

To evaluate the discriminative capacity of the AP-1 signature in predicting BCG response, receiver operating characteristic (ROC) curve analysis was performed using the pROC package (v1.19.0.1)^63^. Patient-level signature scores were used as the predictor variable, with clinical outcome as the response. The directionality of the test was specified such that higher scores corresponded to recurrence. Model performance was quantified using the area under the ROC curve (AUC).

#### External Validation of AP-1 Signature

To validate the AP-1 transcriptional signature in an independent cohort, publicly available single-cell RNA-seq data (GSE269877)^35^ were analyzed. The dataset was imported as a Seurat object, and analysis was restricted to myeloid populations annotated as macrophages and myeloid-derived suppressor cells (MDSCs). Cells were further subset to include only treatment-naïve samples corresponding to groups annotated as “Naive_wo” (responders) and “Naive_w” (non-responders).

Per-cell AP-1 module scores were computed using the same gene set and identical AddModuleScore implementation as described in the discovery cohort to ensure methodological consistency. Patient-level scores were derived by averaging module scores across all cells within each patient, thereby preserving the same aggregation strategy used in the primary analysis.

Patients were grouped according to clinical annotation, with “Naive_wo” classified as responders and “Naive_w” as non-responders. Differences in AP-1 signature activity between groups were assessed using a two-sided Wilcoxon rank-sum test. Gene-level expression of individual AP-1 components was additionally visualized across outcome groups to support interpretation of the composite signature.

#### Bulk RNA-seq Pathway Scoring and Survival Analysis

To evaluate whether immunosenescence–associated inflammatory signalling programs identified in circulating myeloid populations are preserved at the tumour level, pathway activity was assessed in bulk RNA-sequencing data from the BRS cohort^36^. Given the cell type–specific nature of the AP-1 transcriptional signature, downstream inflammatory pathways enriched in association with this phenotype were selected for analysis, including the interferon gamma response, TNF-alpha via NF-κB, complement, and TGF-β signalling gene sets from the Molecular Signatures Database (MSigDB v2026.1.Hs)^29^.

Pathway activity was quantified on a per-sample basis using gene set variation analysis (GSVA)^37^ implemented in the Python package gseapy (v1.1.11)^57^, enabling estimation of coordinated transcriptional programs independent of absolute gene expression levels. Resulting GSVA enrichment scores were used as input for unsupervised clustering to define tumour subgroups.

Hierarchical clustering was performed on GSVA scores using Ward’s linkage as implemented in scikit-learn (v1.8.0)^64^, grouping tumours into distinct phenotypic clusters based on shared pathway activity profiles. Clusters were subsequently annotated based on relative pathway enrichment patterns (*e.g.*, concordant high versus low inflammatory signalling).

Progression-free survival analysis was conducted to evaluate clinical differences between identified clusters. Kaplan–Meier survival curves were generated using the lifelines Python package (v0.30.3)^65^, and statistical significance between groups was assessed using the log-rank test implemented in scipy (v1.17.0)^66^.

## Acknowledgements

This study was supported by grants from the Canadian Institutes of Health Research (Project Grant #173383), the Terry Fox Research Institute (Program Project Grant #1138-05), Marathon of Hope Cancer Centres Network (#3262-02), and the CUASF-BCC Research Grant Program.

