## Supplemental Figures for "Failure of Bacillus Calmette-Guérin therapy in patients with bladder cancer is characterized by immune dysfunction associated with Activator Protein 1"

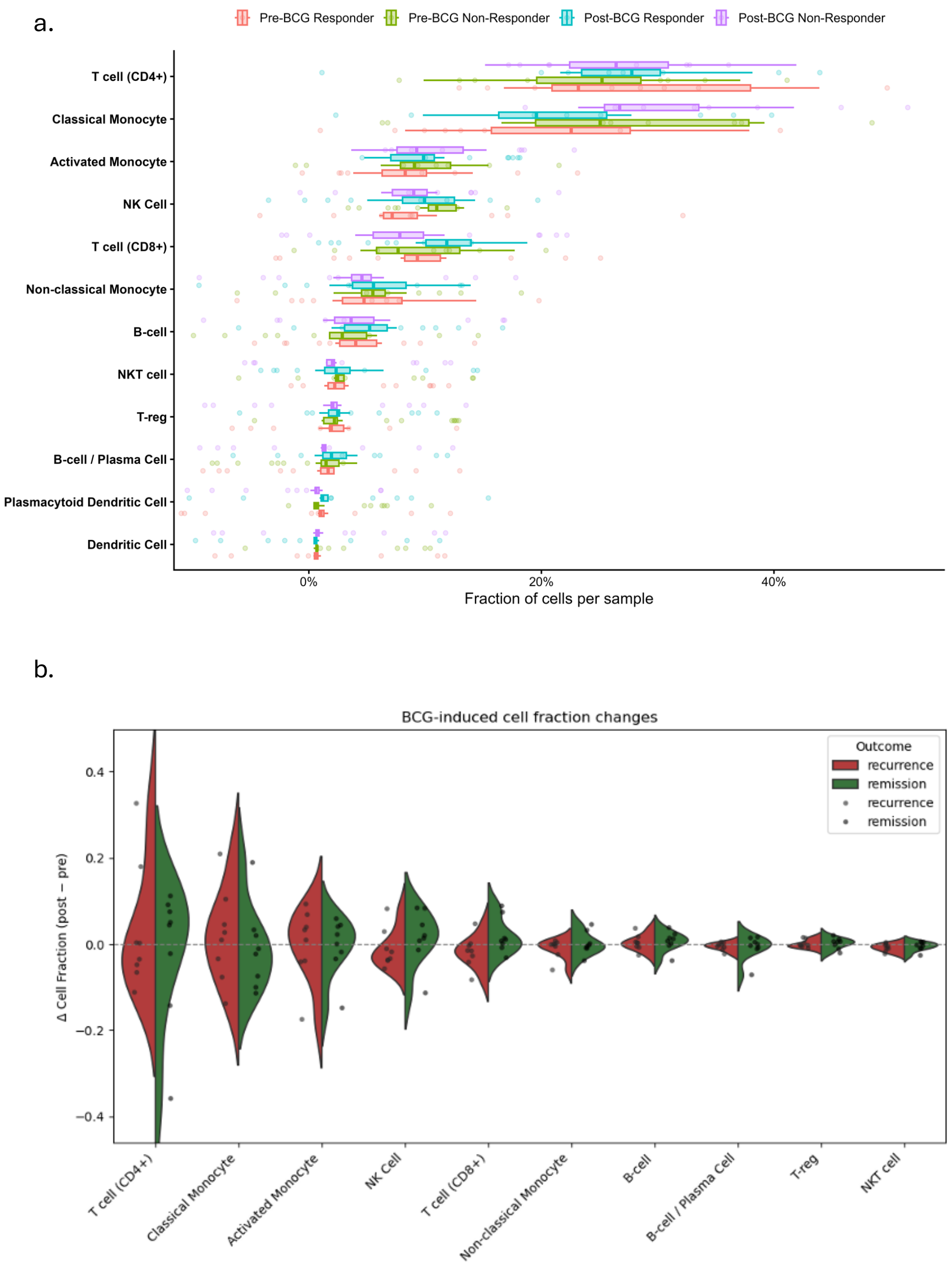


***Supplementary Figure 1. Baseline cellular composition and BCG-induced changes in PBMC cell populations do not differ substantially between responders and non-responders.***

*(a) Distribution of cell type proportions across PBMC samples from BCG responders (remission) and non-responders (recurrence) in both pre- and post-BCG settings. Each point represents the fraction of a given cell type within an individual sample, with boxplots summarizing group-level distributions. Across all major immune compartments, including myeloid (classical, activated, and non-classical monocytes) and lymphoid populations (CD4 $^{+}$T cells, CD8 $^{+}$T cells, NK cells, and B cells), baseline cellular composition was broadly comparable between responders and non-responders, with no significant differences observed between groups.*

*(b) Patient-matched changes in cell type composition following BCG therapy, calculated as the difference in cell fraction between post- and pre-BCG samples (Δ cell fraction). Each point represents an individual patient, with distributions shown separately for responders and non-responders. BCG-induced changes in immune cell composition were modest and largely consistent across response groups, with no major shifts in the relative abundance of key immune populations.*

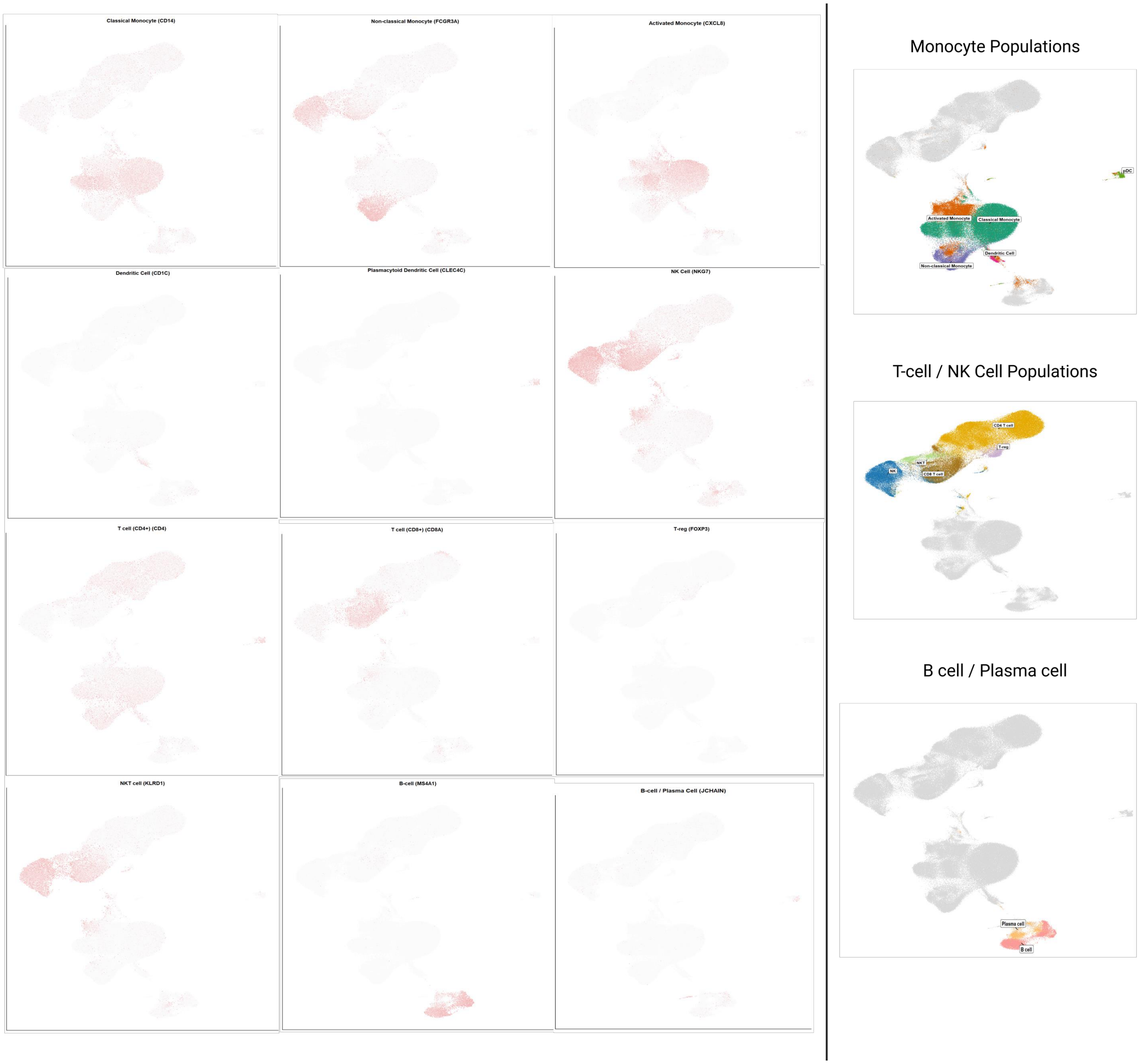


***Supplemental Figure 2. Integrated single-cell transcriptomic landscape of PBMC populations identified in patients undergoing BCG therapy.*** *Uniform manifold approximation and projection (UMAP) visualization of integrated PBMC single-cell RNA sequencing data demonstrating the major immune cell populations identified across all samples. Feature plots display the transcriptomic expression of representative marker genes used to annotate each population, including classical monocytes (CD14), non-classical monocytes (FCGR3A), activated monocytes (CXCL8), dendritic cells (CD1C), plasmacytoid dendritic cells (CLEC4C), NK cells (NKG7), CD4* $^{+}$*T cells (CD4), CD8* $^{+}$*T cells (CD8A), T regulatory cells (FOXP3), NKT cells (KLRD1), B cells (MS4A1), and plasma cells (JCHAIN). Population-level annotation was based on the combined expression of multiple established lineage markers rather than any single transcript alone. Right panels show annotated UMAP embeddings highlighting the major monocyte, T/NK-cell, and B-cell/plasma-cell compartments.*

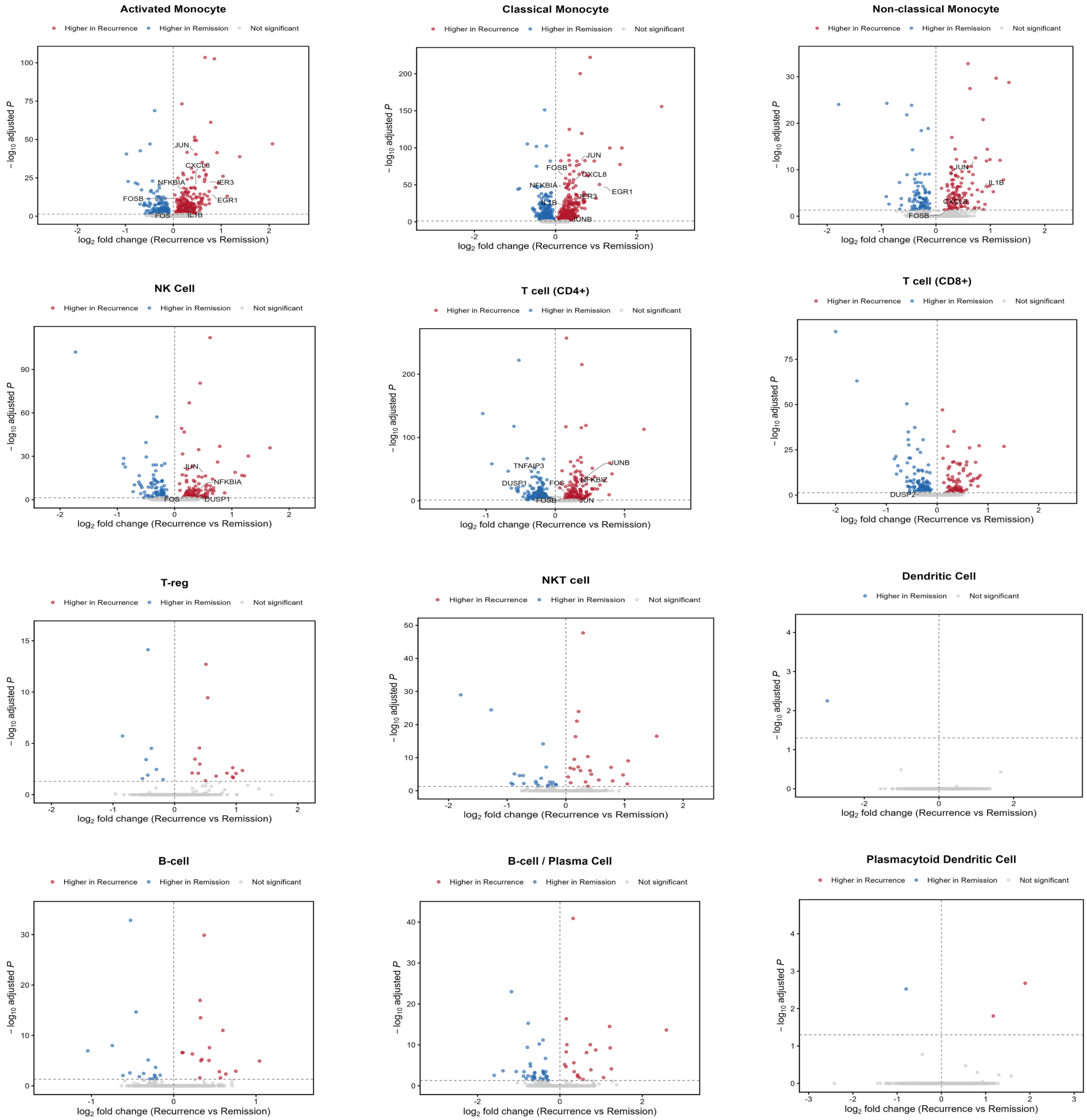


***Supplementary Figure 3. Baseline transcriptional differences between BCG responders and non-responders across PBMC cell types prior to therapy.***

*Volcano plots depicting cell-level Wilcoxon-based differential gene expression analysis comparing BCG responders and non-responders across major PBMC cell types in the pre-BCG setting. Each point represents a gene, plotted by $log_2$ fold change (non-responder vs responder) and $-log_{10}$ adjusted P value. Genes significantly upregulated in non-responders (adjusted P < 0.05) are shown in red, while those upregulated in responders are shown in blue; non-significant genes are shown in grey. Differential expression was most pronounced in monocyte populations, particularly classical and activated monocytes, with comparatively fewer transcriptional differences observed in lymphoid compartments. Select significantly differentially expressed genes associated with pro-inflammatory signalling, NF-κB regulation, AP-1 transcriptional activity, and cellular stress responses (e.g., CXCL8, CCL2, NFKBIA, TNFAIP3, JUNB, FOS, ATF3, IER3, DUSP1, HSPA6) are highlighted.*

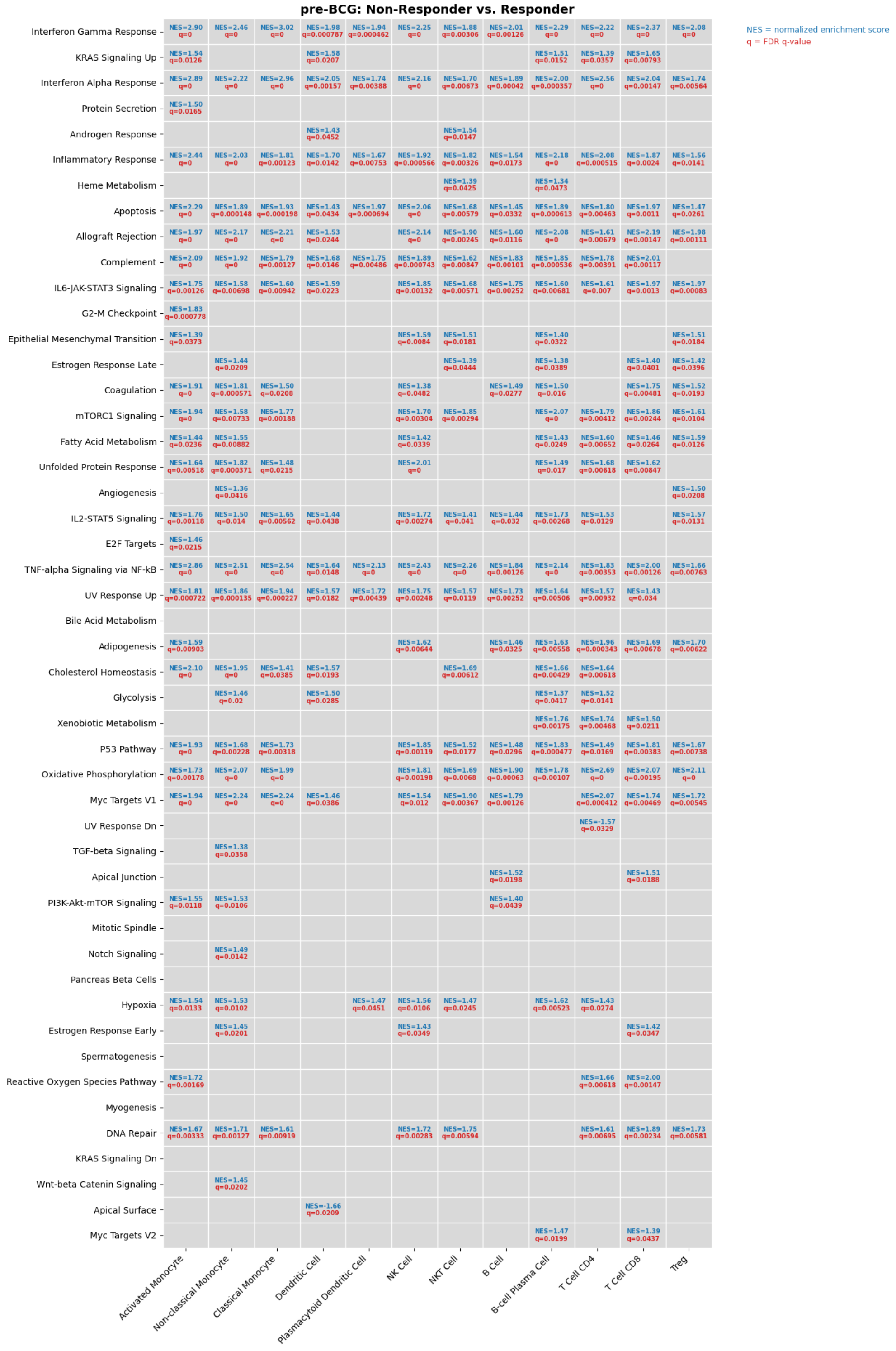


***Supplementary Figure 4. Baseline Hallmark pathway enrichment differences between BCG responders and non-responders prior to therapy.*** *Gene set enrichment analysis (GSEA) was performed across immune cell populations using pre-BCG PBMC samples to compare patients who subsequently responded to BCG therapy versus those who experienced disease recurrence. Rows represent MSigDB Hallmark pathways and columns represent immune cell populations. Significant pathway enrichments (false discovery rate (FDR) q-value < 0.05) are annotated within each tile with the normalized enrichment score (NES; blue text) and associated FDR q-value (red text). Positive NES values indicate relative enrichment in BCG non-responders, whereas negative NES values indicate relative enrichment in BCG responders.*

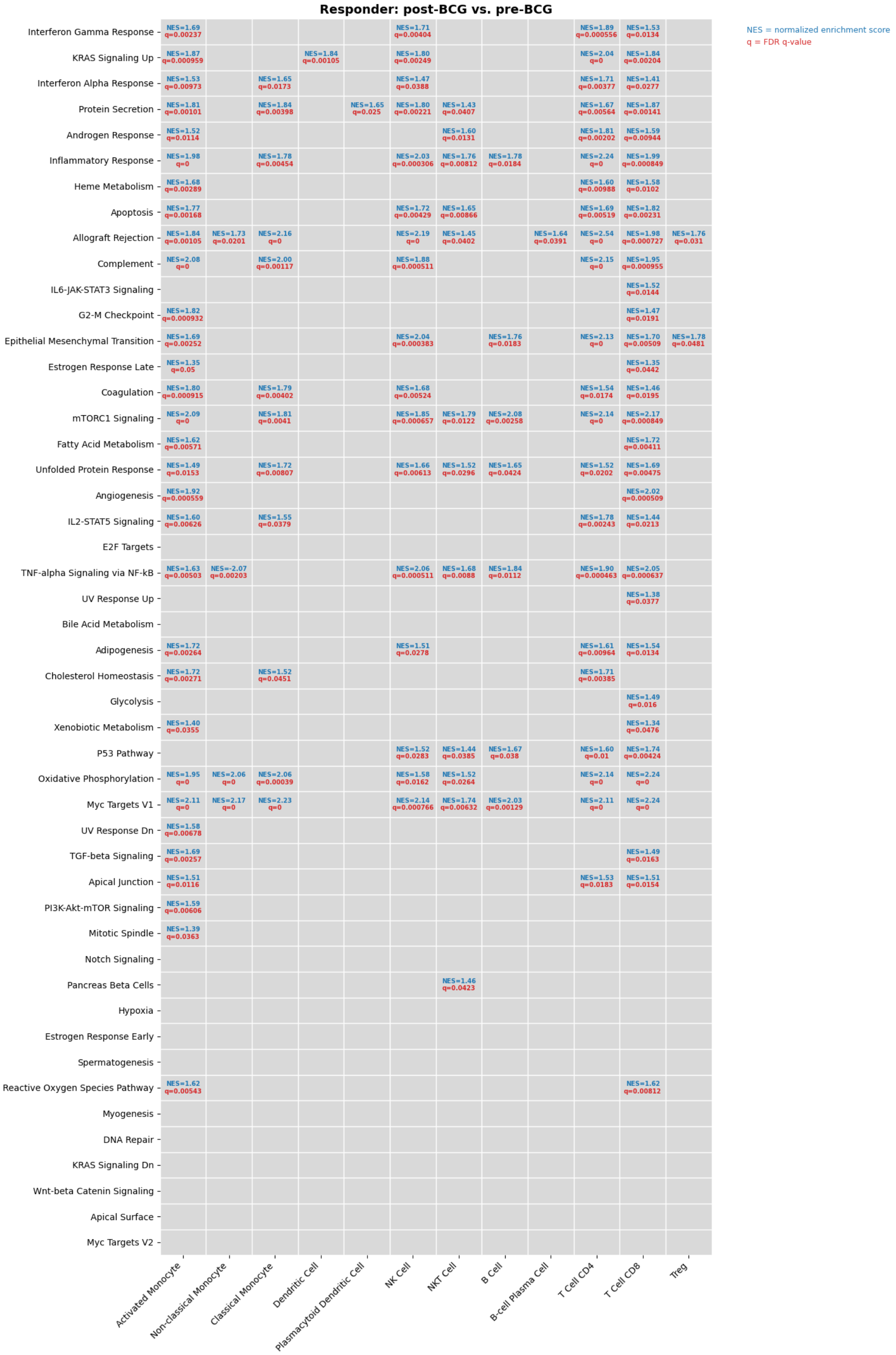


***Supplementary Figure 5. Hallmark pathway enrichment across immune cell populations in BCG responders following therapy.***
*Gene set enrichment analysis (GSEA) was performed across immune cell populations to compare post-BCG versus pre-BCG transcriptional states in patients who remained recurrence-free following BCG therapy (responders). Rows represent MSigDB Hallmark pathways and columns represent immune cell populations. Significant pathway enrichments (false discovery rate (FDR) q-value < 0.05) are annotated within each tile with the normalized enrichment score (NES; blue text) and associated FDR q-value (red text). Positive NES values indicate relative enrichment in post-BCG samples, whereas negative NES values indicate relative enrichment in pre-BCG samples.*

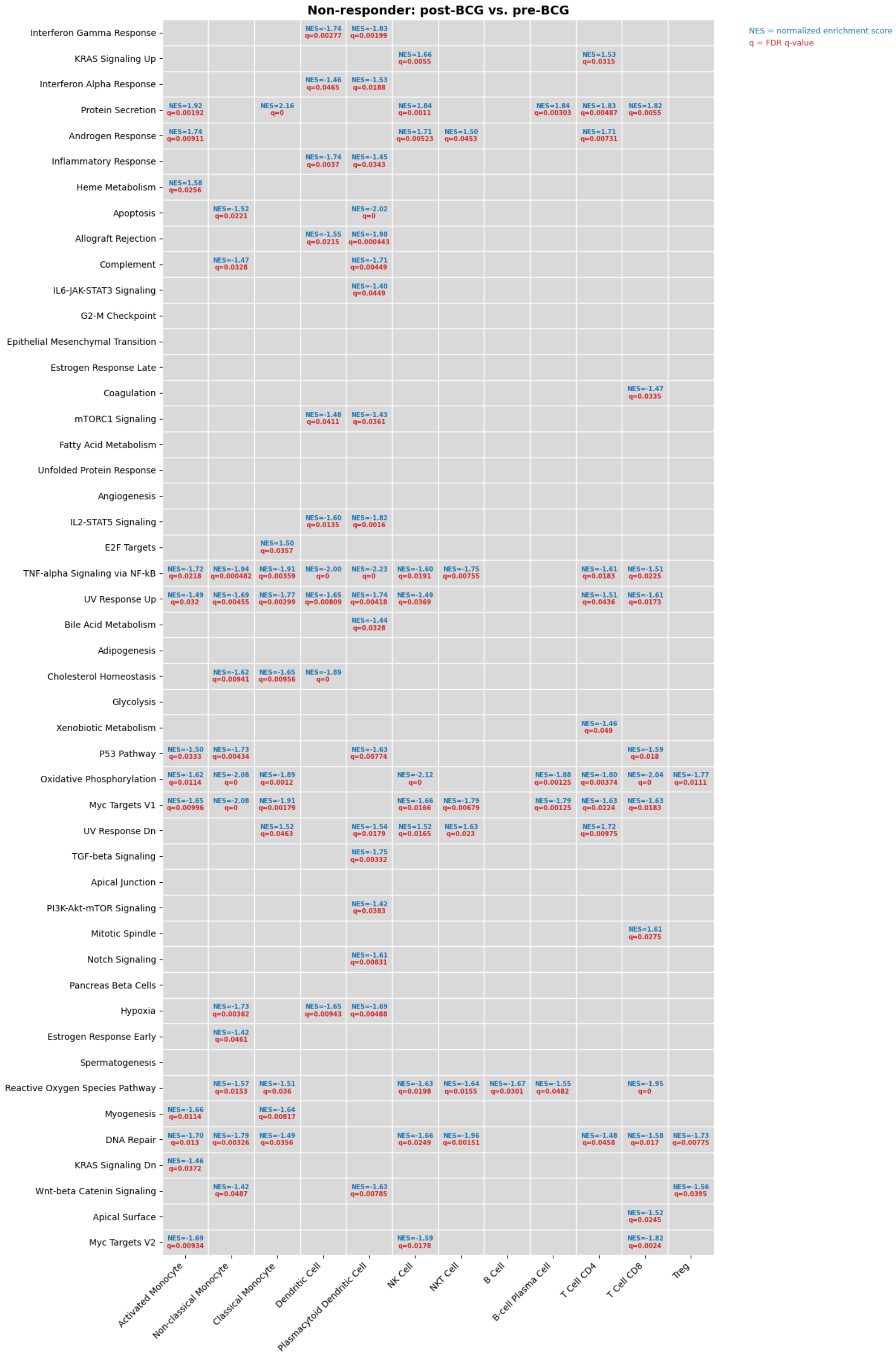


***Supplementary Figure 6. Hallmark pathway enrichment across immune cell populations in BCG non-responders following therapy.***
*Gene set enrichment analysis (GSEA) was performed across immune cell populations to compare post-BCG versus pre-BCG transcriptional states in patients who experienced disease recurrence following BCG therapy (non-responders). Rows represent MSigDB Hallmark pathways and columns represent immune cell populations. Significant pathway enrichments (false discovery rate (FDR) q-value < 0.05) are annotated within each tile with the normalized enrichment score (NES; blue text) and associated FDR q-value (red text). Positive NES values indicate relative enrichment in post-BCG samples, whereas negative NES values indicate relative enrichment in pre-BCG samples.*

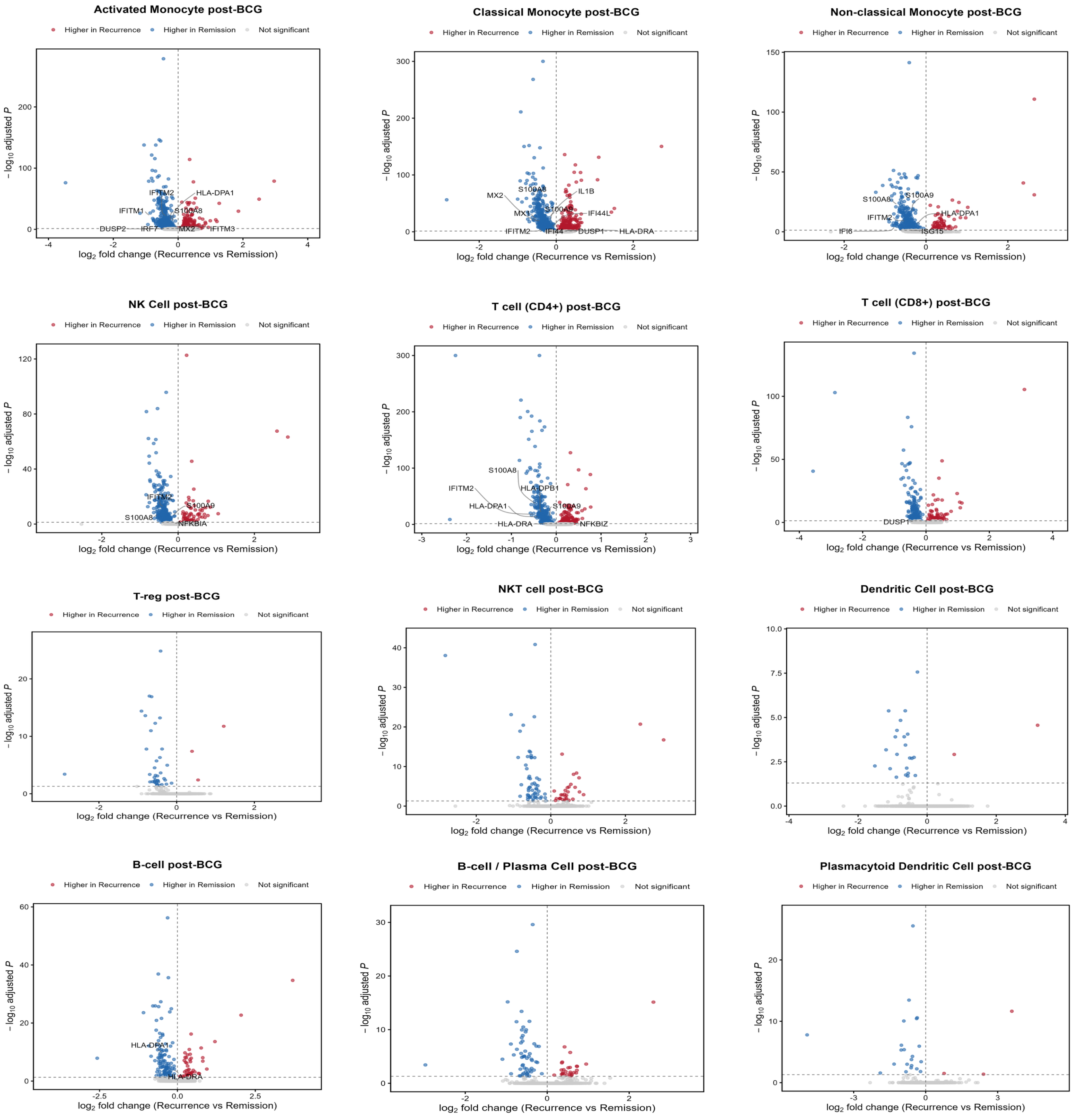


***Supplementary Figure 7. Post-BCG transcriptional differences between responders and non-responders reveal modest but distinct immune activation signatures.***

*Volcano plots depicting cell-level Wilcoxon-based differential gene expression analysis comparing BCG responders and non-responders across major PBMC cell types following BCG induction therapy. Each point represents a gene, plotted by $\log_2$ fold change (non-responder vs responder) and $-\log_{10}$ adjusted P value. Genes significantly upregulated in non-responders (adjusted $P < 0.05$) are shown in red, while those upregulated in responders are shown in blue; non-significant genes are shown in grey. Compared with the pre-treatment setting, transcriptional differences were reduced in magnitude, with the largest number of differentially expressed genes observed in activated and classical monocyte populations, and comparatively fewer differences detected in lymphoid cell types. Genes upregulated in responders were enriched for pro-inflammatory and immune activation programs, including cytokines and chemokines (e.g., CCL20), interferon-stimulated genes (e.g., IFITM1), and immune regulatory factors (e.g., FGL2, IL1B), consistent with transcriptional responses associated with BCG-induced immune activation.*

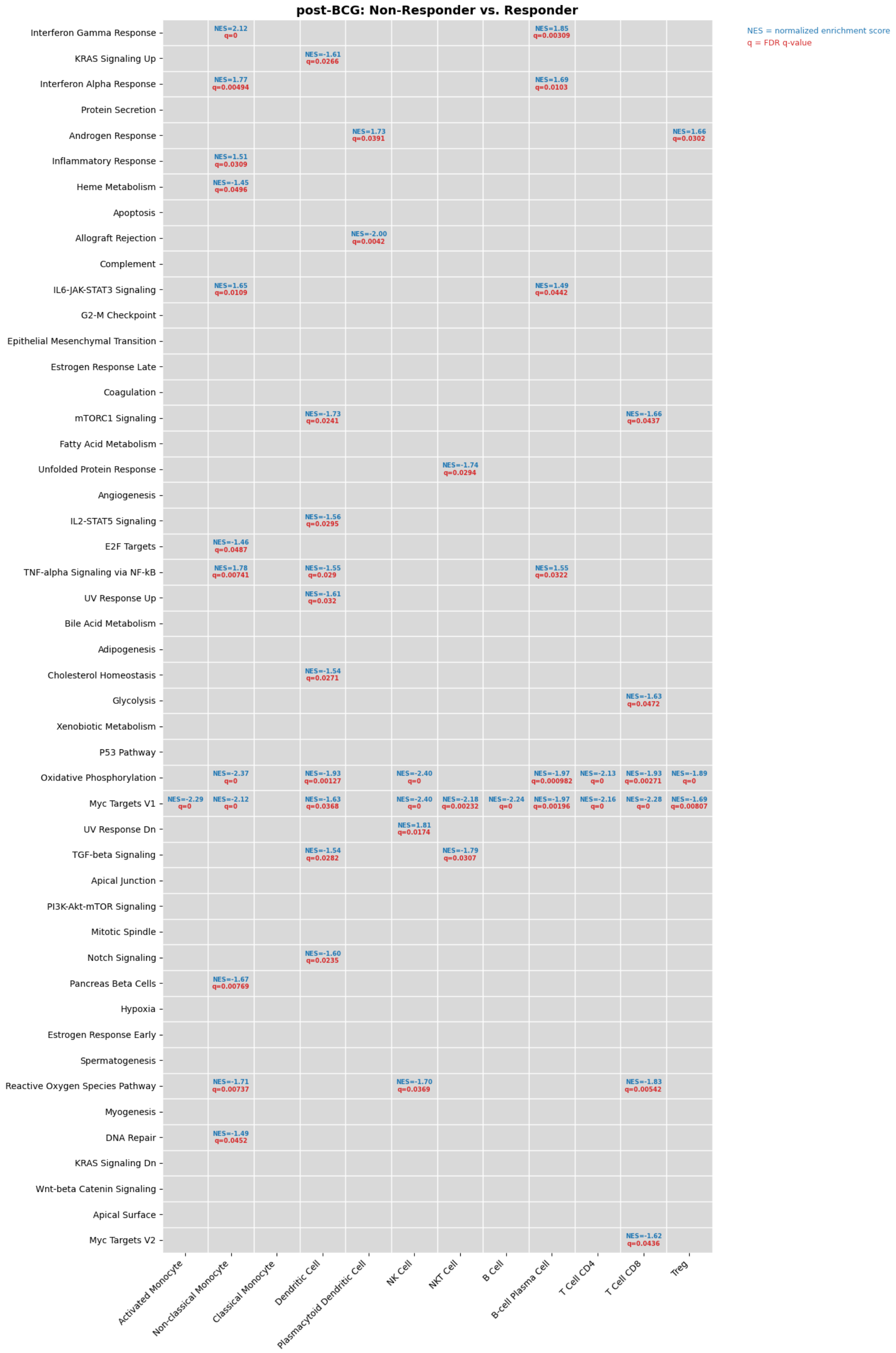


***Supplementary Figure 8. Hallmark pathway enrichment differences between BCG responders and non-responders following therapy.*** *Gene set enrichment analysis (GSEA) was performed across immune cell populations using post-BCG PBMC samples to compare patients who remained recurrence-free following BCG therapy versus those who experienced disease recurrence. Rows represent MSigDB Hallmark pathways and columns represent immune cell populations. Significant pathway enrichments (false discovery rate (FDR) q-value < 0.05) are annotated within each tile with the normalized enrichment score (NES; blue text) and associated FDR q-value (red text). Positive NES values indicate relative enrichment in BCG non-responders, whereas negative NES values indicate relative enrichment in BCG responders.*

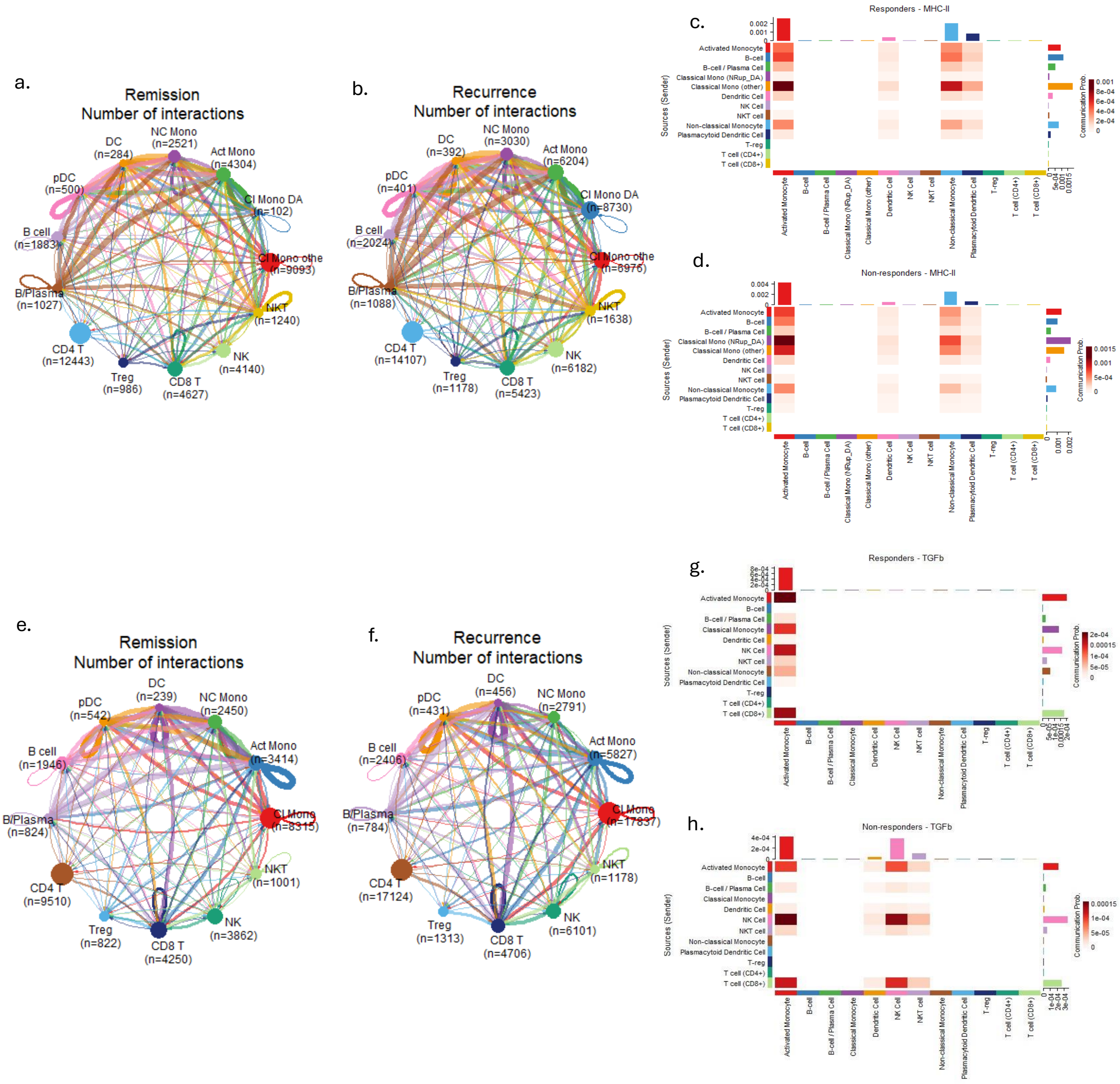


***Supplementary Figure 9. Differential cell–cell communication dynamics distinguish BCG responders and non-responders before and after therapy.***

*Cell–cell communication analysis was performed using CellChat on PBMC single-cell RNA-seq data from BCG responders (remission) and non-responders (recurrence), in both pre- and post-BCG settings. (a–d) Pre-BCG. Circular network plots depict the number of inferred ligand–receptor interactions between immune cell populations in responders (a) and non-responders (b). Node size reflects cell abundance, and edge thickness represents the number of significant interactions. Non-responders exhibit a greater number of inferred interactions, with increased connectivity involving myeloid populations, particularly activated and classical monocytes. Pathway-level communication analysis highlights relative information flow across signalling pathways, demonstrating broadly elevated but diffuse signalling in non-responders (c–d). Notably, non-responders show modestly increased signalling through MHC-II– and TGFβ-associated pathways, alongside reduced CD86-associated co-stimulatory signalling relative to responders. (e–h) Post-BCG. Following therapy, this pattern is reversed. Circular network plots show that responders exhibit a greater number of inferred interactions and enhanced network connectivity involving both myeloid and lymphoid populations (e), whereas non-responders display comparatively reduced connectivity (f). Differential pathway contribution analysis reveals a shift toward increased signalling activity in responders across multiple immune-related pathways, including antigen presentation (MHC-II/MHC-I) and co-stimulatory signalling (g–h). In contrast, signalling in non-responders remains diminished and more restricted across cell types.*

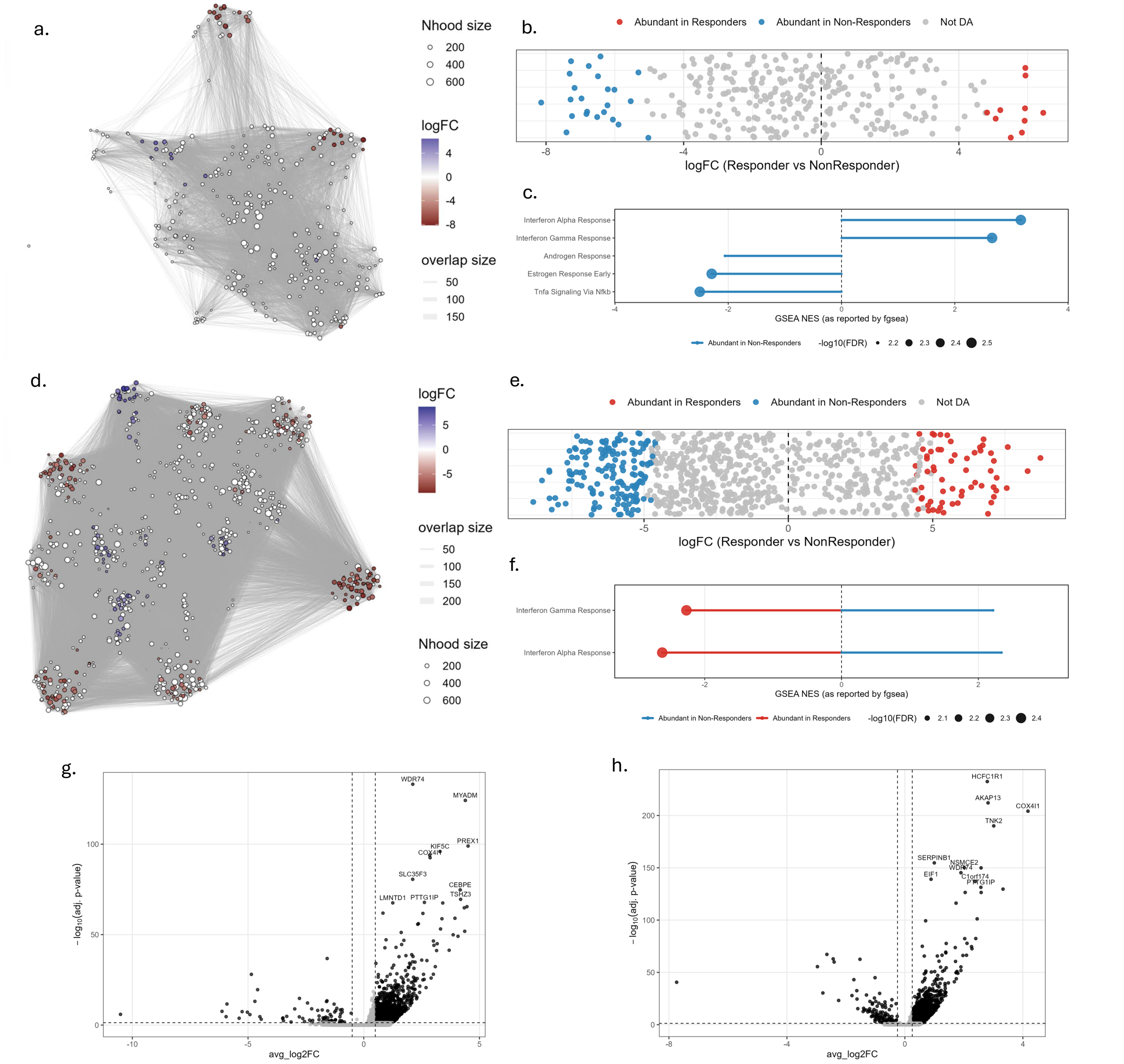


***Supplementary Figure 10. Post-BCG differential abundance and transcriptional programs in monocyte subsets distinguish responders from non-responders following BCG induction therapy.*** *Differential abundance (DA) and regulatory analyses were performed on post-treatment PBMC single-cell multiomic data obtained one week after the fifth induction instillation of BCG, comparing patients who subsequently remained disease-free versus those who experienced recurrence. (a–c) Activated monocyte analyses. (a) Milo-based differential abundance analysis identifies neighbourhoods enriched between groups within activated monocytes. Each node represents a neighbourhood, with colour indicating log fold-change (BCG responders vs BCG non-responders). Blue neighbourhoods denote regions enriched in responders, whereas red neighbourhoods denote regions enriched in non-responders. (b) Distribution of differential abundance statistics across activated monocyte neighbourhoods demonstrates a greater number of neighbourhoods enriched in non-responders following BCG therapy. (c) Gene set enrichment analysis of differentially abundant activated monocyte neighbourhoods reveals increased interferon alpha response and interferon gamma response signalling in responder-enriched neighbourhoods, whereas non-responder-enriched neighbourhoods show relative enrichment of estrogen response, androgen response, and TNFα signalling via NF-κB pathways. (d–f) Equivalent analyses in classical monocytes. (d) Milo-based differential abundance analysis identifies distinct responder- and non-responder-enriched neighbourhoods within classical monocytes following BCG therapy. Blue neighbourhoods indicate enrichment in responders and red neighbourhoods indicate enrichment in non-responders. (e) Differential abundance statistics demonstrate extensive post-treatment remodelling of classical monocyte neighbourhood composition between clinical response groups. (f) Pathway enrichment analysis of differentially abundant classical monocyte neighbourhoods demonstrates increased interferon alpha and interferon gamma response signalling in responder-enriched neighbourhoods, whereas these pathways are relatively diminished in neighbourhoods enriched in non-responders. (g,h) Differential chromatin accessibility analyses of responder-enriched neighbourhoods identify distinct accessible regulatory regions within activated monocytes (g) and classical monocytes (h) following BCG therapy. Volcano plots display differentially accessible peaks between responders and non-responders, highlighting regions associated with genes involved in inflammatory, metabolic, and immune-regulatory programs, including MYADM, PREX1, COX4I1, SERPINE1, and AKAP13, consistent with sustained chromatin remodelling and immune activation in responders after BCG therapy.*

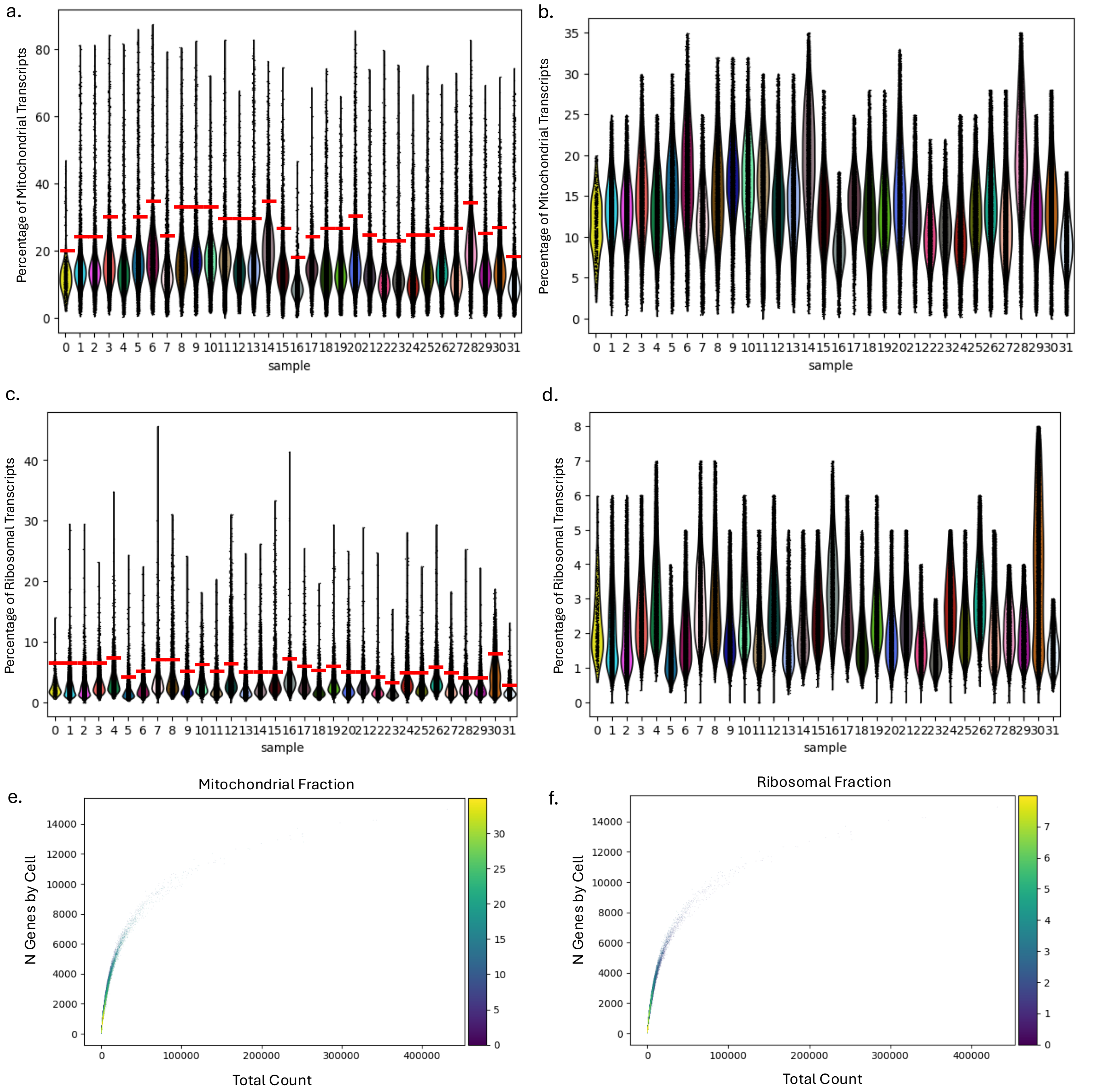


***Supplemental Figure 11. Quality control filtering of single-cell transcriptomic profiles based on mitochondrial and ribosomal transcript fractions.*** *Quality control assessment was performed independently for each sample prior to downstream single-cell RNA sequencing analysis. (a) Distribution of mitochondrial transcript percentages across all cells in each sample before filtering. Red horizontal bars indicate the sample-specific mitochondrial transcript threshold used for quality control filtering. (b) Distribution of mitochondrial transcript percentages following filtering, demonstrating removal of cells with elevated mitochondrial transcript content consistent with low-quality or stressed cells. (c) Distribution of ribosomal transcript percentages across all cells in each sample before filtering, with red horizontal bars indicating the sample-specific ribosomal transcript filtering threshold. (d) Distribution of ribosomal transcript percentages following filtering, demonstrating retention of cells within acceptable ribosomal transcript ranges. (e,f) Scatter plots showing the relationship between total transcript counts (UMIs) and the number of detected genes per cell following filtering, coloured by mitochondrial transcript fraction (e) and ribosomal transcript fraction (f), respectively. These plots illustrate the distribution of retained high-quality cells across sequencing depth and transcriptional complexity following quality control filtering.*

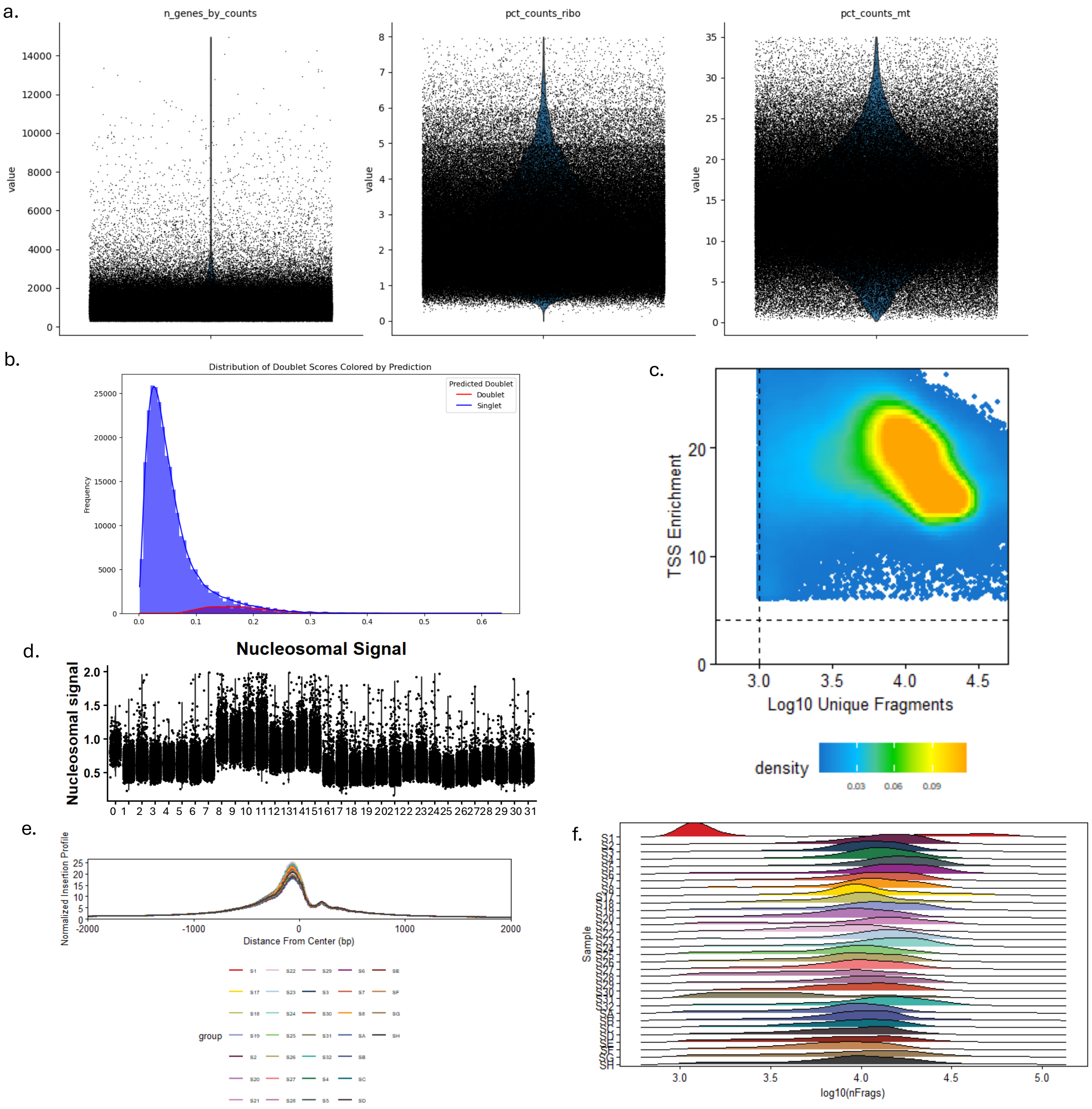


***Supplemental Figure 12. Quality control assessment and filtering of single-cell transcriptomic and chromatin accessibility profiles.*** *Quality control metrics were evaluated prior to downstream single-cell multiomic analysis to identify low-quality cells, technical artifacts, and doublets. (a) Distributions of the number of detected genes per cell (n_genes_by_counts), ribosomal transcript fraction (pct_counts_ribo), and mitochondrial transcript fraction (pct_counts_mt) across all cells following initial filtering. These metrics were used to assess transcriptional complexity and cellular integrity. (b) Distribution of Scrublet-derived doublet scores coloured by predicted classification, showing separation between predicted singlets and putative doublets. Cells exceeding the inferred doublet threshold were removed from downstream analyses. (c) Density plot of scATAC-seq quality metrics showing transcription start site (TSS) enrichment versus log10 unique fragments per nucleus. Dashed lines indicate the minimum thresholds applied for retention of high-quality nuclei based on chromatin accessibility complexity and TSS enrichment. (d) Distribution of nucleosomal signal values across all samples. Nucleosomal signal, defined as the ratio of mononucleosomal to nucleosome-free fragments, was used to identify low-quality nuclei, and nuclei with nucleosomal signal values greater than 2 were excluded from downstream analysis. (e) Aggregate transcription start site enrichment profiles across all samples, demonstrating strong enrichment of accessible chromatin signal at TSS regions consistent with high-quality ATAC-seq libraries. (f) Distribution of log10 unique fragments per nucleus across samples, illustrating chromatin accessibility complexity and consistency in sequencing depth between samples following quality control filtering.*
